# Inhalable point-of-care urinary diagnostic platform

**DOI:** 10.1101/2023.09.30.560328

**Authors:** Qian Zhong, Edward K.W. Tan, Carmen Martin-Alonso, Tiziana Parisi, Liangliang Hao, Jesse D. Kirkpatrick, Tarek Fadel, Heather E. Fleming, Tyler Jacks, Sangeeta N. Bhatia

**Author notes:** These authors contributed equally to this work (Qian Zhong, Edward K.W. Tan).

## Abstract

The late-stage detection of lung cancer leads to a high global mortality rate. Although low-dose computed tomography screening improves lung cancer survival in at-risk groups, this test still suffers from high rates of false positive results. In addition, inequality remains in the diagnosis of lung cancer as access to medical imaging infrastructure is limited. Here, we designed a needleless and imaging-free platform, termed PATROL (**<u>p</u>**oint-of-care **<u>a</u>**erosolizable nanosensors with **t**umor-**<u>r</u>**esponsive **<u>ol</u>**igonucleotide barcodes), to increase detection accuracy, to reduce resource disparities for early detection of lung cancer, and to enable timely interception. PATROL formulates a set of DNA-barcoded, activity-based nanosensors (ABNs) into inhalable formats that can be delivered using clinical nebulizers or inhalers. Lung cancer-associated proteases in the tumor microenvironment selectively cleave the ABNs, releasing synthetic DNA reporters that are eventually excreted via the urine. The barcoded nanosensor signatures present in urine samples are quantified within 20 minutes using a multiplexable paper-based lateral flow assay at room temperature. PATROL detects early-stage tumors in an autochthonous lung adenocarcinoma mouse model with high sensitivity and specificity. Tailoring the library of ABNs may enable the modular PATROL platform to not only lower the resource thresholds required for early detection of lung cancer, but also enable rapid detection of chronic pulmonary disorders and infections.

## Main

Lung cancer death has continued to decline in nations with a high development index (HDI), due in part to significant progress in early detection tools and timely therapeutic interception.^1^ However, disproportionately high mortality is observed in lung cancer cases in low- and middle-income countries (LMICs) and correlates with late-stage disease detection, illustrating inequity in early diagnosis in resource-poor settings— one of the chief challenges in addressing cancer health disparities.^2^ Where available, low-dose computed tomography (LDCT) has been the standard of care to screen for early-stage lung cancer among high-risk and asymptomatic people, and this practice has led to an approximately 20-25% reduction of mortality in clinical trials.^3–5^ And yet, reduced patient access to imaging and the scarcity of trained personnel remain significant clinical challenges in areas outside urban imaging infrastructures in HDI countries, not to mention in LMICs.

Liquid biopsies (LBs) that detect the presence of endogenous, cancer-implicated biomarkers in body fluids, such as shed proteins, circulating tumor DNA (ctDNA), and tumor cells, are emerging as transformative, noninvasive tools to diagnose and monitor lung cancer.^6–9^ These assays often rely on resource-intensive analytical technologies such as chromatography, tandem mass spectrometry, next-generation sequencing, and flow cytometry to identify molecular or genetic changes present in captured samples, thus limiting their applicability in low-resource settings. Furthermore, the detection of cancer-associated biomarkers is conditional upon their release into the bloodstream, which places constraints on other activity-based hallmarks of diseases from being utilized for diagnostic purposes. For instance, proteases are found to play direct functional roles during tumor progression, and display distinct expression and activity signatures across hallmarks of cancer, but are physically located in the tumor microenvironment (TME), and thus might be overlooked by LBs.^10^ We and others have developed exogenous biomarkers—activity-based diagnostics (ABDs)—that leverage catalytic proteases to continuously liberate synthetic barcodes. These catalytically-amplified and measurable exogenous reporters reveal the signature of proteolytic dysregulation in the TME that are linked to cancer states and therapeutic responses via imaging, urine biopsy, or breath biopsy,^11–18^ thereby expanding the panel of available analytes for more predictive diagnostics. We envision that the development of point-of-care (POC) tests that offer accurate real-time ABD readouts could further unleash their translational potential and alleviate surging demands for early lung cancer detection tools and improve cost-effectiveness of this screening practice, especially in resource-poor settings. Lateral flow assays (LFAs), a well-established platform for POC testing, have been utilized to help assess at-risk groups by detecting the presence of cancer-associated proteins and genetic mutations in LBs.^19,20^ However, cancer-detecting LFAs have been minimally implemented, in part due to the need for multiplexing capacity, and the multistep assay protocols necessary to derive quantitative measurements. Inspired by the multiplexed nature and potential for immobilization of nucleic acids on paper strips, the rational design of chemically-stabilized oligonucleotides for molecular barcoding might enable accurate LFA-based readouts of dysregulated proteolysis through fluid biopsy.

In addition to reducing the resource demands of detection assays, diagnostic platforms can also be improved by establishing self-administered formulations, as this would further lower the threshold for clinical deployment. For example, inhaled medicines have been routine tools for at-home treatment of chronic lung diseases for decades.^21,22^ Indeed, this foundation has been leveraged during the design of cutting-edge inhalation technologies (e.g. nebulization, dry powder inhalers) to achieve pulmonary delivery of diagnostics, allowing them to bypass non-specific systemic degradation and profile early lesions in a proximal manner.^15,23,24^ Aerodynamic size often dictates the deposition of aerosols in human respiratory tracts and lungs. Larger particles are intercepted by adsorption in the mouth, throat, and upper airways, whereas smaller particles (i.e. 1-5 µm) are more likely to deposit in the peripheral lung (bronchi, bronchioles, or alveoli) where lung cancer is predominantly located. Particles <0.5 μm are typically exhaled, and as such, deep-lung deposition of these very small particles is reduced.^25^ Therefore, the complexity of nanoparticle delivery via inhalation arises from the need to quickly convert nanoscale formulations to aerosol clouds with (aerodynamic) size neither too small nor too big — ideally 1-3 μm in diameter.

Here, we introduce a highly modular platform for early detection of lung cancer in a POC format. PATROL integrates 3 modules — low-plex activity-based nanosensors (ABNs), a portable inhalation unit, and a multiplexable paper-based LFA (**Figure 1)**. To enable precision diagnostics, we combined transcriptomic data from human samples with insight from an earlier, intratracheal diagnostic study in mice,^23^ and nominated protease substrates specific to Stage I lung adenocarcinoma (LUAD) to probe for tumor-associated proteolytic signatures. We re-engineered DNA-barcoded ABNs into micrometer-sized aerosol formulations to optimize the potential for deposition in human lungs. The inhalable ABN format exhibits excellent aerodynamic performance and offers considerable promise for noninvasive, self-administered human delivery by employing clinical nebulizers or hand-held inhalers. In a genetically-engineered mouse model of LUAD, after individual mice breathed in the nebulized aerosols, ABNs were delivered uniformly to lungs, where the substrates exposed to the TME were selectively cleaved to shed synthetic DNA barcodes into the systemic circulation. The nucleic acid reporters encoding unique tumor proteolytic signatures were subsequently concentrated in the urine via the kidney. For rapid detection of the DNA barcodes at the POC, we engineered LFAs to quantify the multiplexed urinary reporters on a single strip at room temperature, and achieved diagnostic results within 20 minutes. Through the integration of different technological components, we have established a non-invasive ‘inhale and detect’ approach to accurately diagnose early-stage lung cancer that would not require trained medical personnel, a long duration treatment, or centralized diagnostic laboratories. Furthermore, the high modularity of PATROL enables its potential to be extended to achieve rapid detection of chronic pulmonary disorders and infections.

**Figure 1.**
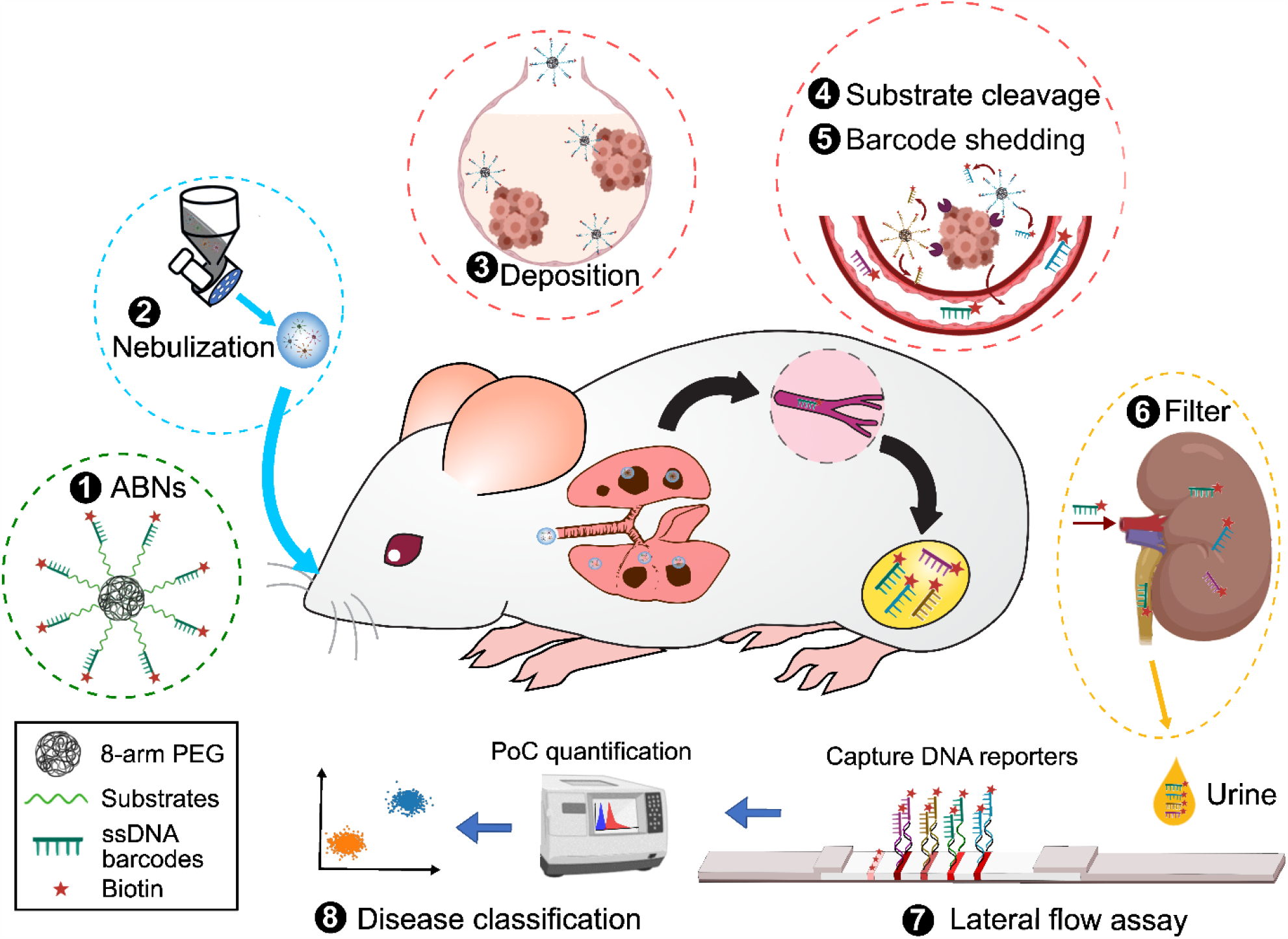
PATROL for early detection of LUAD at the point-of-care. **1)** Single-stranded, phosphorothioate-modified DNA (ssDNA) 20mers and protease substrates were conjugated to 8-arm PEG nanoscaffolds via click chemistry, creating a cohort of multiplexed ABNs with synthetic DNA barcodes. The DNA barcodes were biotinylated, which can later bind Europium-neutravidin complex on paper strips of an LFA. **2)** Non-invasive pulmonary delivery of DNA-barcoded ABNs via inhalation. Atomization of nanoscale ABNs into micro-sized aerosols (nano-in-micro process) using a vibrating-mesh nebulizer enabled efficient respiratory deposition of inhaled ABNs. The resulting aerosols were introduced into a nose-only inhalation tower with airflow. **3)** The aerosols were spontaneously inhaled by mice with early-stage autochthonous LUAD^*26*^ and deposited on their airways and lung periphery where LUAD nodules reside. **4)** The DNA barcodes were quickly liberated upon substrate cleavage by dysregulated proteases associated with early LUAD, and **5)** were shed into the systemic circulation. **6)** These distinct DNA barcodes in circulation become concentrated via the kidney into the urine. **7)** The urine was sampled 120 minutes after ABN inhalation and a customized paper-based LFA with multiplexing capacity quantified DNA barcode concentrations present in urine samples at room temperature in a POC manner. The LFA is based on the sequence-specific hybridization at room temperature and labelling of designated DNA barcodes at each of the test lines. The fluorescence of each signal line present on the LFA was quantified using a POC reader, and **8**) disease classification was carried out based on principal component analysis.

## Results

### Engineering ABN formulations for inhalable diagnostics

As particles with 1-5 µm in aerodynamic diameter tend to deposit in human peripheral lungs (**Fig. 2A**),^27^ we first sought to re-engineer ABNs into submicron aerosols, so as to be suitable for delivery via inhalation (**Fig. 2B**) via nebulizers. We began by synthesizing a conventional ABN by functionalizing an 8-arm polyethylene glycol (PEG) nanoscaffold with tandem peptides containing an a matrix metalloproteinase (MMP0 substrate (i.e., LQ81 in **Table S1**) and a Cy7-tagged Glu-fibrinopeptide B (GluFib) reporter. Each PEG scaffold carried approximately 8 tandem peptides and the hydrodynamic size ranged from 10 to 15 nm in diameter. We chose nebulizers for the proof of concept as they require minimal formulation design to generate aerosols, and inhalers were further benchmarked against nebulizers. The model ABNs were dissolved in 0.9% NaCl solution at a range of concentrations up to 2 mM substrate equivalent (or 250 µM PEG equivalent) and then aerosolized with a vibrating mesh nebulizer. We observed that aerosol production by the nebulizer was considerably reduced when the substrate concentration exceeded 1 mM (or 0.125 mM PEG equivalent), and so proceeded with ABN formulations incorporating no more than 1 mM substrates. Nebulizers create aerosols continuously, while aerosols are breathed in only during inspiratory phases. We next sought to quantify the inhaled proportions of ABN aerosols, defined as delivered doses (DDs). To measure DDs in vitro, a simplified breath simulator was used to create representative profiles (i.e., sinusoidal) of human respiration and the aerosols were collected during inhaling cycles in a next generation impactor (NGI) (**Fig. 2B, Fig. S1A**) – a simplified model of human tracheobronchial tree and alveoli. The DDs of aerosols generated from various ABN concentrations remained largely unchanged at 35-38% (**Fig. 2C**) and the residual volume of condensed aerosols in the nebulizer and T tubing was similar and minimal (data not shown). NGI collects aerosol samples in stages as they pass through a cascade of progressively finer nozzles (**Fig. 2B**,**D**), and the mass of aerosols deposited in each stage can be used to estimate aerodynamic size and fine particle dose (**Fig. 2D**). The median mass aerodynamic diameters (MMADs) of ABN aerosols generated by the vibrating mesh nebulizer were 3.72-4.35 µm (**Fig. 2D, E**) and the size change was inversely proportional to the ABN concentration. Conversely, the fine particle fractions (FPFs) of these aerosols, a parameter that estimates aerosol fractions that favor peripheral lung depositions, were proportional to the ABN concentrations, culminating at 60.2% of the DDs (**Fig. 2F**). Both DDs and FPFs fall within the range of inhalation drugs tested with the same nebulizer. The size and proteolytic sensitivity of the ABNs before and after nebulization were largely unaltered in comparison with the original nanosensors (**Fig. S1B, Fig. 2G**).

**Figure 2.**
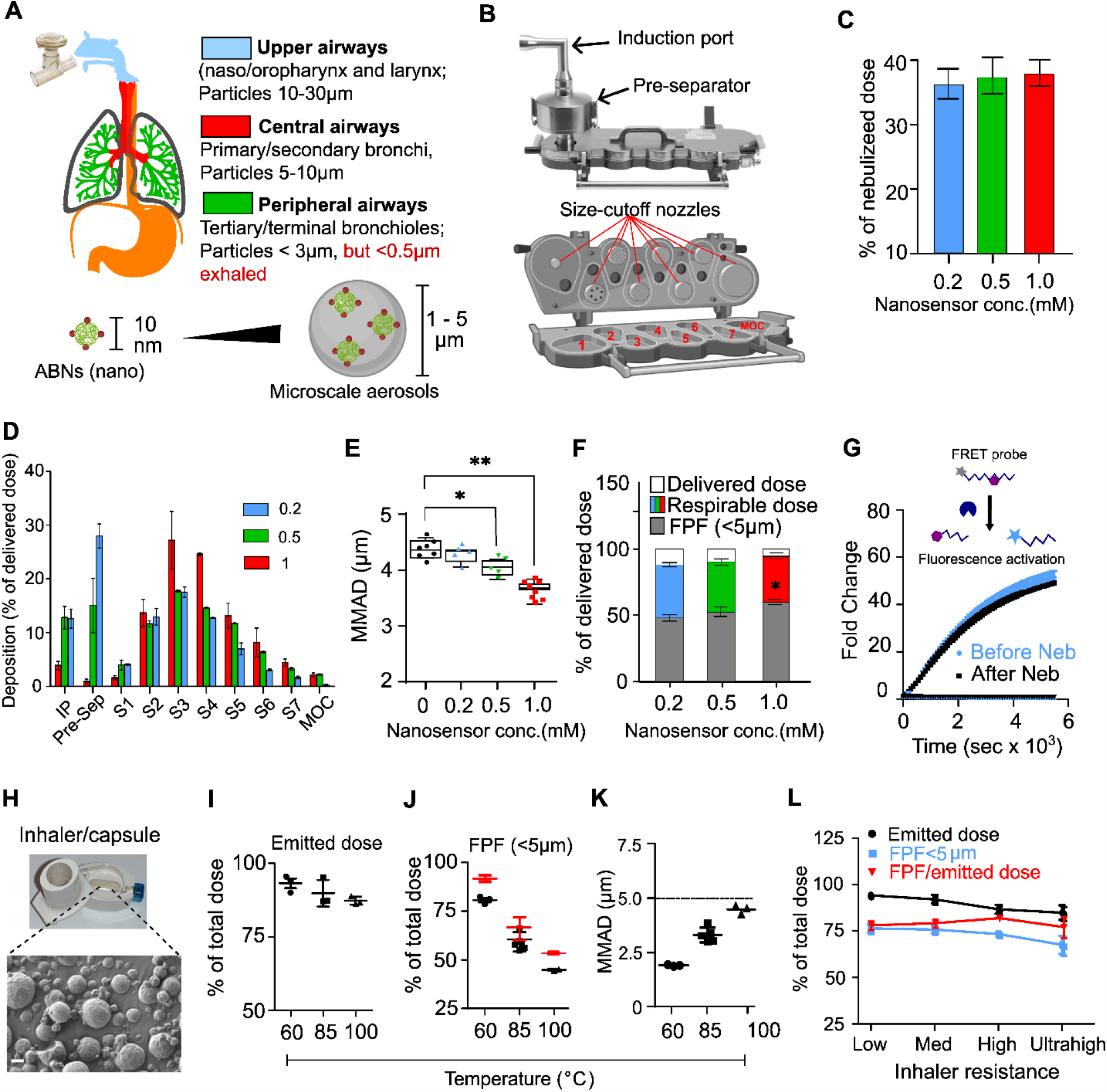
Engineering inhalable formulations of ABNs to noninvasively probe protease dysregulation. **(A)** Human adult airways and lung geometry, with predicted aerosol deposition with respect to particle size. Efficient deposition of ABNs on peripheral lungs requires nanoscale ABNs to be formulated into aerosols with a preferred aerodynamic size of 1-5 microns in diameter. The aerosols can be generated via nebulizers or inhalers. **(B)** Next generation impactor (NGI) and a transverse section, or inside view (below). The NGI classifies aerosols into known size range by drawing the aerosols through a cascade of progressively finer nozzles (pointed to by red lines). The air jets from these nozzles impact on plane sampling surfaces of Stage 1-7 and MOC, and each stage collects finer aerosols than its predecessor. Number 1-7 = Stage 1-7, MOC=micro-orifice. **(C)** Delivered dose (DD) was minimally changed with the concentration of ABNs in bulk solution. The DD represents a fraction of total nebulized aerosols that are inhaled by subjects under a standardized breathing pattern of 500 mL tidal volume, 1:1 inhalation:exhalation (I:E) ratio, and 15 breaths per minute (BPM) frequency, which imitates human breathing cycle – inspiration and expiration. **(D)** Aerosol deposition (by mass) of the nebulized ABNs on Stage 1-7 and MOC was measured with the setup in **Fig. S2A** under the standardized breathing profile described in (C). IP=induction port, Pre-Sep=pre-separator, S1-7= Stage 1 to 7, and MOC= micro-orifice. The 0.2, 0.5 and 1 mM denote the concentration of peptide substrates in the bulk solutions used for nebulization, which also correspond to 25, 62.5, and 125 μM of 8-arm PEG nanoscaffolds, as each PEG nanoscaffold carried ∼8 peptides. **(E)** The median mass aerodynamic diameter (MMAD) of ABN aerosols generated by a vibrating-mesh nebulizer ranged from 3.72 to 4.35 µm, with a downward trend as the concentration of ABNs in bulk solution rose. The concentration of nanosensors is equal to that of peptide substrates. **(F)** Fine particle fractions (FPFs, grey) of nebulized aerosols under a continuous air flow of 15 L/min increased as the ABN concentration increased. FPFs were calculated with particle sizing data acquired with a next generation impactor. FPF is defined as the dose fraction of inhaled particles with aerodynamic size less than 5 μm in diameter, representing the amount of cargo that can potentially reach the lungs. The detailed calculation of delivered dose in (C), MMAD in (D), and FPF in (F) is provided in **Supplementary Information**. *p < 0.05, and **p < 0.01 by One-way ANOVA test with Dunnett’s multiple comparisons to the size of 0 µM PEG nanoscaffold in (E) and (F), respectively. **(G)** The relative release of a quenched fluorescent reporter after incubation with recombinant MMP13 was measured for ABNs before and after nebulization. Reconstituted ABNs from nebulized aerosols demonstrated similar cleavage susceptibility to the recombinant MMP13 as the original, un-nebulized nanoparticles. FRET-paired peptides turn on fluorescence when MMP13 cleaves the substrate and liberates the quencher. **(H)** Scanning electron microscopic image of ABN-laden microparticles. Microparticles were spray-dried at 60 °C. Scale bar = 1 µm. **(I)** Emitted doses, **(J)** FPFs, and **(K)** MMADs of ABN-containing microparticles calculated with aerosols collected in the stages of NGI. The microparticles were spray-dried at different inlet temperatures – 60, 85, and 100°C. 15 mg microparticles were loaded into a capsule and tested with a monodose RS01 inhaler. Emitted dose is a proportion of microparticles discharged from an inhaler with respect to total dose. Red dots in (J) represent FPFs recalculated by dividing FPFs by emitted doses. **(L)** Emitted dose and FPF are largely independent of inhaler intrinsic resistance. Statistical analysis was performed with respect to the low-resistance inhaler using one-way ANOVA with Dunnett’s multiple comparisons and no statistical significance was found. To reach 4 KPa pressure drop over the inhaler, low-, medium-, high-, or ultrahigh-resistance inhalers require air flow rate = 90L/min, 68 L/min, 53L/min, or 33L/min in NGI, respectively.

To enable high dose delivery to deep lungs and improve cost effectiveness and ease to use of inhalation devices for point-of-care detection, we also explore the feasibility of formulating the nanosensors into portable, electric power-free, and hand-held devices — dry powder inhalers (DPIs) that are capable of delivering large doses of submicron powders (dry aerosols) to the lungs within seconds upon a single breath actuation. We utilized a particle engineering technology, termed excipient enhanced growth (EEG), to make carrier-free and micrometer-sized dry particles (1-3μm) that increase in size following inhalation in order to minimize upper respiratory tract deposition and maximize targeted deposition in the lung periphery.^28^ A solution of nanosensor cocktails, mannitol (a hygroscopic excipient) and L-leucine (a surface-active dispersion enhancer) were co-sprayed via a spray-dryer (**Fig. S1C**) where the solution is heated at the inlet and sprayed through fine nozzles into a dryer, followed by exit from the outlet in the form of submicron powders. The resulting microparticles show relatively coarse surface morphology (**Fig. 2H**). These dry particles can release the nanosensors immediately after reconstitution in simulated lung fluids (**Fig. S1D**), indicating the potential for fast dissolution of the inhaled microparticles when in contact with humid respiratory tract. We then loaded these microparticles into a capsule and inserted it in a clinically-used inhaler -RS01. A short period of air flow (approx. 3-4 s and contingent on inhaler resistance) mimicking a deep inhalation disaggregates and introduces microparticles into NGI to assess aerodynamic performance. In the preliminary optimization, we found that inlet temperature (heating the incoming ABN/D-mannitol/L-leucine solution) dictates aerodynamic performance of the inhalable microparticles in comparison with other parameters such as feed flow rate, outlet temperature, and air flow rate. The spray-drying at inlet temperature = 60°C yields the highest emitted dose (92.3±3.0% of total mass of microparticles loaded in the capsule, **Fig. 2I**) and FPF (91%±1.3% of emitted dose, **Fig. 2J**), and the lowest MMADs (2.1±0.1μm, **Fig. 2K**). This suggests that nearly all microparticles not only are respirable, but also can potentially reach the deep lungs (i.e. regions rich in bronchioles and alveoli). We then assessed the aerodynamic performance of ABN-laden microparticles in dry powder inhalers with different intrinsic resistance (low, medium, high and ultrahigh). Despite emitted doses and FPFs decrease slightly as inhaler resistance (**Fig. 2L**), all the inhalers can discharge more than 80% microparticles (emitted dose) and generated 78-91% fine particles (of emitted dose) conducive to peripheral lung deposition. Although the low-resistance inhaler seems the best performer in the assessment, high inspiratory airflow rates and respiratory efforts required by such inhalers cannot always be achieved by high-risk or elderly populations suffering chronic pulmonary disorders. The aerosol performance is largely independent of inhalers, indicating highly comparable delivery efficiency and dose uniformity of the inhalable, ABN-containing microparticles across different inhalers. Collectively, these protease nanosensors can be formulated for efficient lung delivery via aerosol through direct nebulization from aqueous solutions, or incorporation into respirable dry microparticles.

### Nominating a panel of ABNs with sufficient power and POC adaptability

To date, heterogenous malignancies are typically classified via high throughput screening of large panels of biomarkers, yet this process requires resource-intensive techniques. However, the design of a POC diagnostic tool requires both facile administration and a low-infrastructure readout. Thus, we sought to identify a minimal set of precision probes that could offer high predictive power via simply carried out screening operations suitable for decentralized settings, and thereby help inform early detection and initiate timely interception. By leveraging tumor proteases, we established a library of ABNs that can detect early-stage lung cancer by building on our previously-validated probes and nominating additional proteases and substrates specific to early lesions,^23^ with the compatibility for low-plex POC assay requirements as our priority. We used DESeq2 to identify differentially-expressed proteases in tumors and non-tumor adjacent tissues (NATs) from patients with Stage I LUAD (**Fig. S2A**) in the Cancer Genome Atlas (TCGA) database. We found significant protease dysregulation even at the earliest stages of disease, with the top 22 upregulated proteases listed in **Fig. S2A**. We then constructed a library of 73 peptide substrates mined from the literature and paired them with quenched fluorescent labels to form activatable probes for *in vitro* screening against the selected, dysregulated proteases (**Fig. S2B**). The probes were incubated with recombinant proteases and we measured the cleavage kinetics as reflected by the fold change in fluorescence signals (illustrated in **Fig. S2B**,**C**). Amongst the 73 nominated substrates, we selected 15 substrates for their selectivity and sensitivity to target proteases (**Fig. 3A**,**B**). We further selected another 5 peptide sequences from our existing substrate library to augment the capacity to probe the activity of metalloproteases and proteases secreted by infiltrating immune cells.^15,16,23^ From the *in vitro* screening, the substrates show high specificity against the family of proteases and preferential selectivity against proteases of the same subtype (**Fig. 3B, Fig. S2B**).

**Figure 3.**
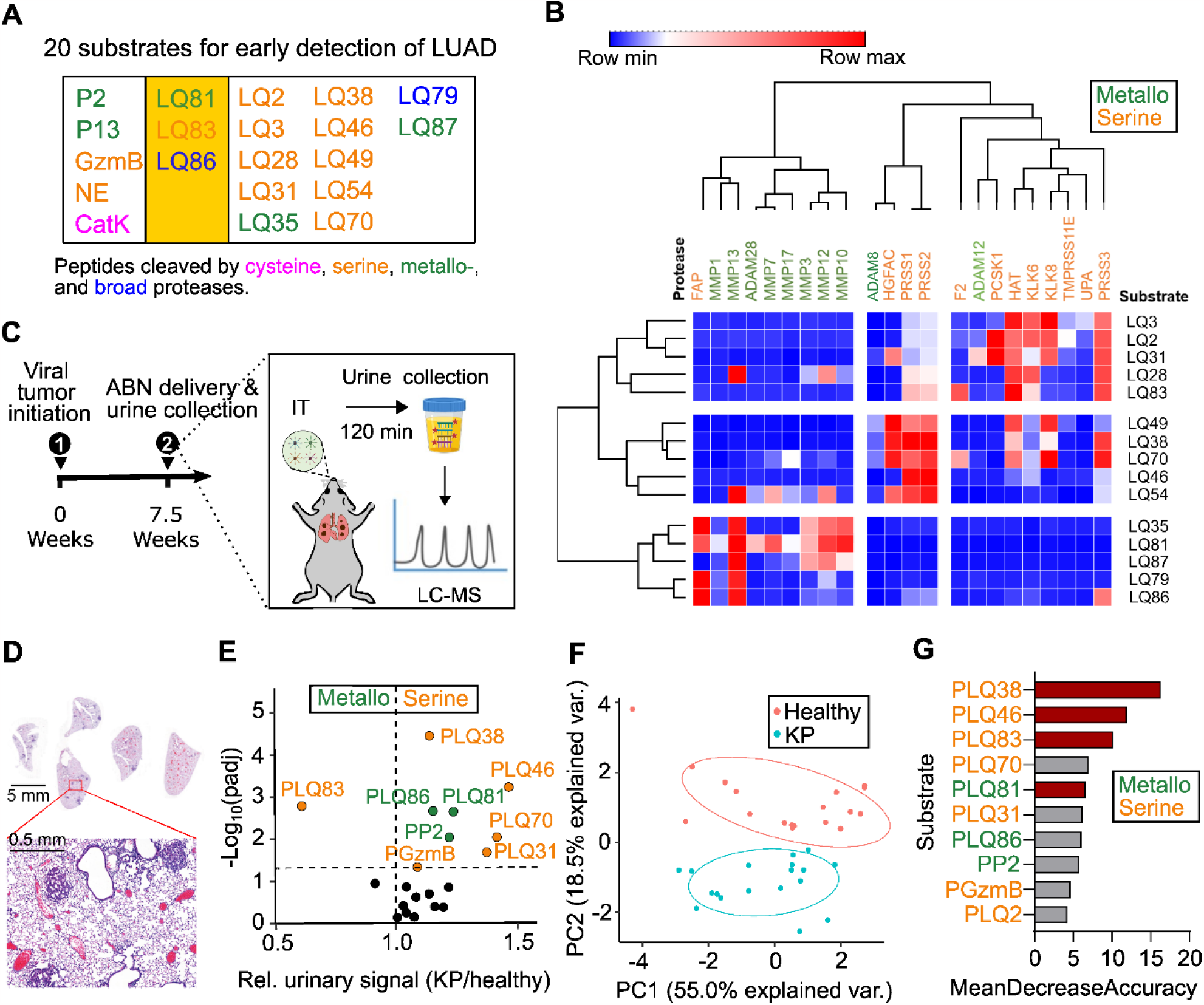
Nomination of protease substrates specific to early-stage lung cancer. **(A)** A panel of 20 peptides (**Table S1**) susceptible to proteolytic cleavage by dysregulated proteases in Stage I LUAD. Substrates in magenta, green, orange and blue are susceptible to cysteine proteases, metallo-proteases, serine proteases, or broadly cleaved by both metallo-/serine proteases, respectively. The peptide substrates in two left-most columns have been previously validated as prominent probes in the classification of LUAD in mouse models^*23*^ and characterizing proteolytic activity of infiltrated myeloid cells, whereas the substrates tabulated in Column 2-5 were exclusively nominated for Stage I human LUAD. Three peptides (2^nd^ yellow column) were included in both groups. **(B)** Heatmap of z-scored fluorescent fold-changes at 50min (average of 2 replicates) showing hierarchical clustering of proteases dysregulated in Stage I LUAD (horizontal) by their substrate specificities and of selected FRET-paired synthetic substrates (vertical) by their protease specificities. The heatmap of 15 substrates (vertical) by 21 proteases (horizontal) was re-tabulated from **Fig. S4C**. Green and orange denote metallo-and serine proteases, respectively. **(C)** Mass-coded ABNs were delivered into KP mice via intratracheal instillation 7.5 weeks after tumor induction. The urine samples were collected at 120 minutes post administration and the mass-coded reporters were quantified with liquid chromatography-mass spectrometry. **(D)** Hematoxylin and eosin staining of lung sections shows KP lung nodules at 7.5 weeks (KP_7.5wk_) after induction were mainly Grade I/II tumors and nodule size was less than 0.5 mm. **(E)** *In vivo* screen of the panel of 20 ABNs shown in (A) revealed that 9 reporters were differentially enriched in the urine of healthy and KP_7.5wk_ mice (green and orange dots). Volcano plot showing statistical significance and fold change of urinary reporters in KP_7.5wk_ mice over healthy counterparts. Each dot represents an ABN probe as labeled. **(F)** Principal component analysis enabled the successful classification of KP mice with early LUAD from the healthy cohort using 20-plex ABNs. Each dot represents one mouse in the animal cohort. **(G)** Variable importance analysis ranked the importance of the probes for tumor classification. Probes with higher mean decrease accuracy (i.e. importance scores) produced by the random forest model, contribute more to the diagnostic performance. Mean decrease accuracy scores express accuracy the model losses by excluding each variable. Four ABNs (PLQ38, PLQ46, PLQ83 and PLQ81) are among the probes with the most robust diagnostic power (red columns), and are collectively susceptible to a range of protease classes (both metallo- and serine). This 4-plex was then nominated for subsequent formulation as an inhalable ABN panel.

The final expanded panel of 20 candidates was then evaluated via *in vivo* screening in mouse models in an effort to nominate a small, bespoke probe set with low-plex compatibility (**Fig. 3A, Table S1**). Specifically, the set of 20-plex ABNs was synthesized by conjugating mass-coded tandem peptides (GluFib isotope+ substrate peptide) to the PEG nanoscaffolds (**Table S2**). We then evaluated the ABNs in an autochthonous LUAD mouse model that was induced in the Kras^LSL-G12D;^Trp53^fl/fl^ (KP) mice by adenovirus expressing Cre recombinase under the control of the surfactant protein C promoter.^26,29^ Previous work established this model as closely recapitulating human disease progression from alveolar adenomatous hyperplasia (AAH) to grade IV adenocarcinoma.^23,26^ The lung tumors were allowed to grow for 7.5 weeks (KP_7.5wk_) at which point lung nodules were histologically categorized as Grade I/II and the nodule sizes were typically smaller than 0.5 mm in diameter (**Fig. 3D**). The pooled 20 ABNs were intratracheally instilled to the KP_7.5wk_ and age/gender-matched healthy mice, and the urine produced 60-120 min after ABN administration was collected. The mass-encoded reporters present in the urine samples were analyzed using tandem liquid chromatography-mass spectrometry/mass spectrometry (LC-MS/MS). As shown in the volcano plot, 9 of 20 probes offer significant detection power (**Fig. 3E**). Unsupervised dimensionality reduction by principal component analysis (PCA), an algorithm that transforms a large set of variables into a smaller one without losing any important information, also showed that this 20-plex ABNs was able to cluster tumor-bearing mice at an early time point (**Fig. 3F**). Using variable importance analysis with the random forest model, we further identified the 4 probes (PLQ38, PLQ46, PLQ81 and PLQ83) among those with the greatest fold change (KP/healthy mice) and significance (**Fig. 3E**,**G**). Notably, PLQ38, PLQ46 and PLQ83 are cleaved by serine proteases such as PRSS, HGFAC, or F2, whereas PLQ81 is a metalloprotease substrate. We thus selected this subset of 4 ABNs to reformulate into a low-plex, inhalable panel and assess its diagnostic performance *in vivo* to detect early-stage LUADs.

### Nebulization of ABNs differentiates early LUAD in autochthonous mouse models

We next sought to validate the inhalable ABN formulations *in vivo* for lung cancer detection. While DPI-based microparticles demonstrate superior potential for deep lung deposition in humans, nebulizers remain a primary candidate for *in vivo* validation given the lack of DPIs designed for rodents (DPIs are breath-actuated), and thus the DPI advantage regarding lung deposition patterns cannot be recapitulated in rodent models. As illustrated in **Fig. S3A**,**B**, mice were placed in restrainers of an inhalation tower and they were exposed to nebulized nanosensors only via nasal openings. We first maximized the delivery efficiency of the aerosols by varying flow rates of air influx into the tower which, in combination with nebulization rates, determined delivered doses. Micropumps that drew air at 25 mL/min mimicked mouse inhaling at three randomly selected ports and Cy7-tagged model ABNs (i.e. PEG-LQ81-GluFib) were delivered via aerosols.^30^ We found that in comparison with 1, 4, 8, and 20 L/min, the flow rate at 2L/min deposited the highest portion of aerosols to each nasal opening, which was approximately 0.6% of the total dose entering the tower (**Fig. S4A**) over 10 minutes of nebulization. Higher flow rates led to more accumulated aerosols in the exhaust filter, while lower flow rates (e.g. 1 L/min) increased residual volume of ABN solutions in the nebulizer and T-tubing hose. We further observed that the nebulized ABNs resulted in a more even distribution throughout the lung periphery, while intratracheally-instilled ABNs tended to accumulate in bronchi and central lungs, with high variability across mice in the tested cohort (**Fig. 4A, Fig. S4B**,**C**). This high heterogeneity of distribution may overwhelm specific respiratory zones but disregard other areas of the tract, leading to the possibility of missing tumor nodules. We also found the concentration of released reporters in plasma and urine was higher in the nebulization group (**Fig. S4D**,**E**) than that from the intratracheal group when dosed with the same amount of ABNs. Differing from intratracheal instillation, inhaled ABNs that deposit on upper airways were cleared into the gastrointestinal system (**Fig. S5A**,**B**) due to mucus entrapment and mucociliary escalation clearance.^13^ To assess whether these ABNs re-enter systemic circulation, we orally gavaged the mice with the ABNs and examined pharmacokinetics, biodistribution, and urinary readouts. We found that no fluorescent signals from either cleaved reporters or intact ABNs were detected outside of the digestive organs (**Fig. S5C**,**D**), suggesting that the swallowed nanosensors would not yield background signals in the urine that lower diagnostic power. The ABNs in the stomach and intestines were excreted within 24 hours (**Fig. S5E**).

**Figure 4.**
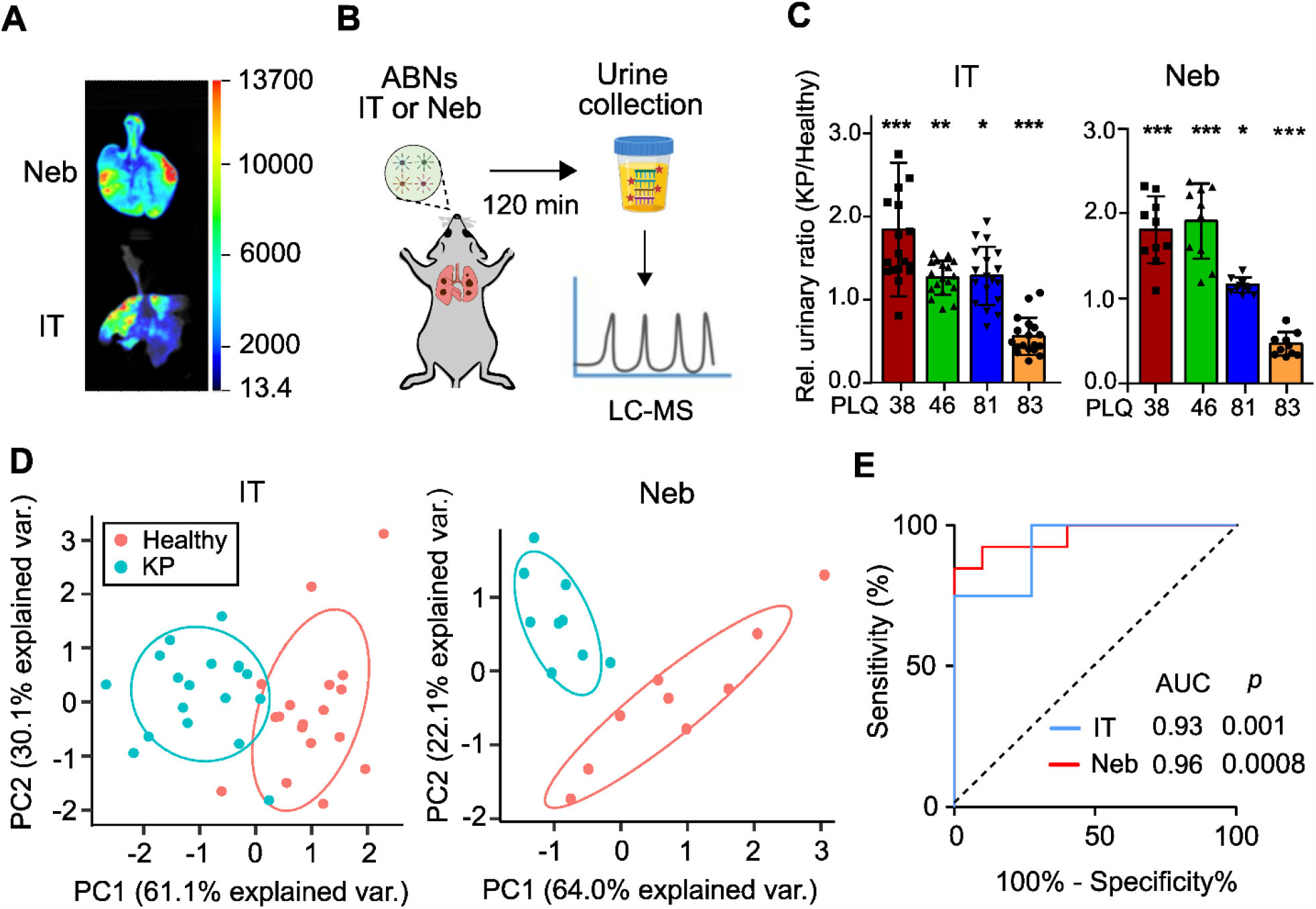
Inhalable ABNs enabled classification of early LUAD. **(A)** Lung local biodistribution of 1 nmol 8-arm PEG nanoscaffolds delivered to wild-type C57B/L via nebulization (Neb, top) or intratracheal instillation (IT, bottom), as characterized by fluorescence imaging. The PEG scaffold was labeled with near infrared dye, Cy7. **(B)** Mass-coded ABNs were delivered into KP mice via either intratracheal instillation or nebulization 7.5 weeks after tumor induction. The urine samples were collected at 120 minutes post administration and the mass-coded reporters were quantified with LC-MS/MS. **(C)** Urinary output of 4-plexed mass-coded ABNs intratracheally delivered (KP: n = 17; healthy control: n = 18) or nebulized (KP: n = 9; healthy control: n = 9) to KP (KP_7.5wk_) and control mice at 7.5 weeks after tumor induction; *p < 0.05, **p < 0.01 and ***p<0.001 by Student’s t-test. Error bars represent SDs. **(D)** Principal component analysis of mean-normalized urinary mass reporters for KP_7.5wk_ and control mice treated with either intratracheally delivered ABNs or with nebulized aerosols. Each dot represents a mouse in the animal cohort. **(E)** ROC curves showing a predictive power of 4-plex urinary reporters from a subset of KP_7.5wk_ and healthy controls in discriminating an independent test cohort of KP lung tumor from healthy controls. A baseline area under the curve (AUC) value of 0.5 represents a random classifier (black dotted line), and a perfect AUC is 1.0. P value represents the significance level between the AUC of KP ROC and that of baseline ROC. Abbreviation: IT=intratracheal, Neb=nebulization.

We next examined whether nebulized 4-plex nanosensors can detect the LUAD at early stages and are used as an alternative to intratracheal instillation. The mass-coded 4-plex ABNs were administered via a nebulizer to the KP_7.5wk_ and healthy mice (**Fig. 4B**). Urinary signals of all 4 reporters were significantly different when administered via nebulizer (PLQ38 and PLQ83: P_adj_<0.001, PLQ46: P_adj_<0.01, and PLQ81: P_adj_<0.05), indicating significant alterations of proteolytic cleavage signature between KP_7.5wk_ and healthy mice (**Fig. 4C**). In comparing the diagnostic capacity of the same 4-plex of sensors administered by the two different routes, we observed that unsupervised dimensionality reduction by PCA was able to classify most of the KP_7.5wk_ mice from healthy counterparts in the group of intratracheal instillation and all KP_7.5wk_ mice in the nebulization group (**Fig. 4D**). We also performed receiver operating characteristic (ROC) analysis, such that the area under the ROC curve (AUC) is calculated as measure of classification accuracy, where a perfect diagnostic has an AUC of 1 and a random diagnostic has an AUC of 0.5. ROC analysis also revealed that nebulized nanosensors (AUC_neb_ = 0.961) performed statistically equivalent to the intratracheally-instilled down-selected (AUC_IT_ = 0.932) library (**Fig. 4E**). With 100% specificity, the nebulized 4-plex ABNs exhibited sensitivity of 84.6%, slightly greater than 75.5% from the intratracheal instillation group but without statistical difference. In summary, inhalable low-level multiplexing of rationally selected ABNs demonstrate robust power for the early detection of mouse autochthonous LUAD.

### Engineered LFA enables room temperature multiplexing of synthetic DNA barcodes

To enable the wider deployment of ABN technology in resource limited settings, we sought to transition to POC detection from the use of peptide-based mass barcodes that require LC-MS/MS for analysis in centralized laboratories. Recently, we reported the use of single-stranded DNA (ssDNA) as barcodes for ABNs.^31^ Here, we sought to design POC assays to quantify a urinary ssDNA barcode signature, with 1) ease of multiplexing, 2) room temperature operation, and 3) high sensitivity and selectivity. In **Fig. 5A**, we outline the design of a multiplexable LFA that operates based on DNA hybridization on a single strip. The target barcodes are 20 nucleotides in length on phosphorothioate backbones with biotin on the 5’ ends for tagging purposes. On the LFA, urinary ssDNA barcodes are first tagged with Eu-labeled neutravidin which is pre-loaded on the conjugate pad. Eu is used as a fluorescent readout for quantitative measurements. The running buffer then carries the Eu-tagged DNA barcodes along the test strip, where the DNA barcodes are subsequently hybridized with their capture sequences, thus forming fluorescing test lines. We also employ a positive control line of biotinylated BSA, such that in the absence of a fluorescent signal at this position, a faulty test result would be inferred. Each test takes about 20 minutes at room temperature and the test strips can then be analyzed by a commercial POC fluorescent reader (Axxin AX-2X-S)

**Figure 5.**
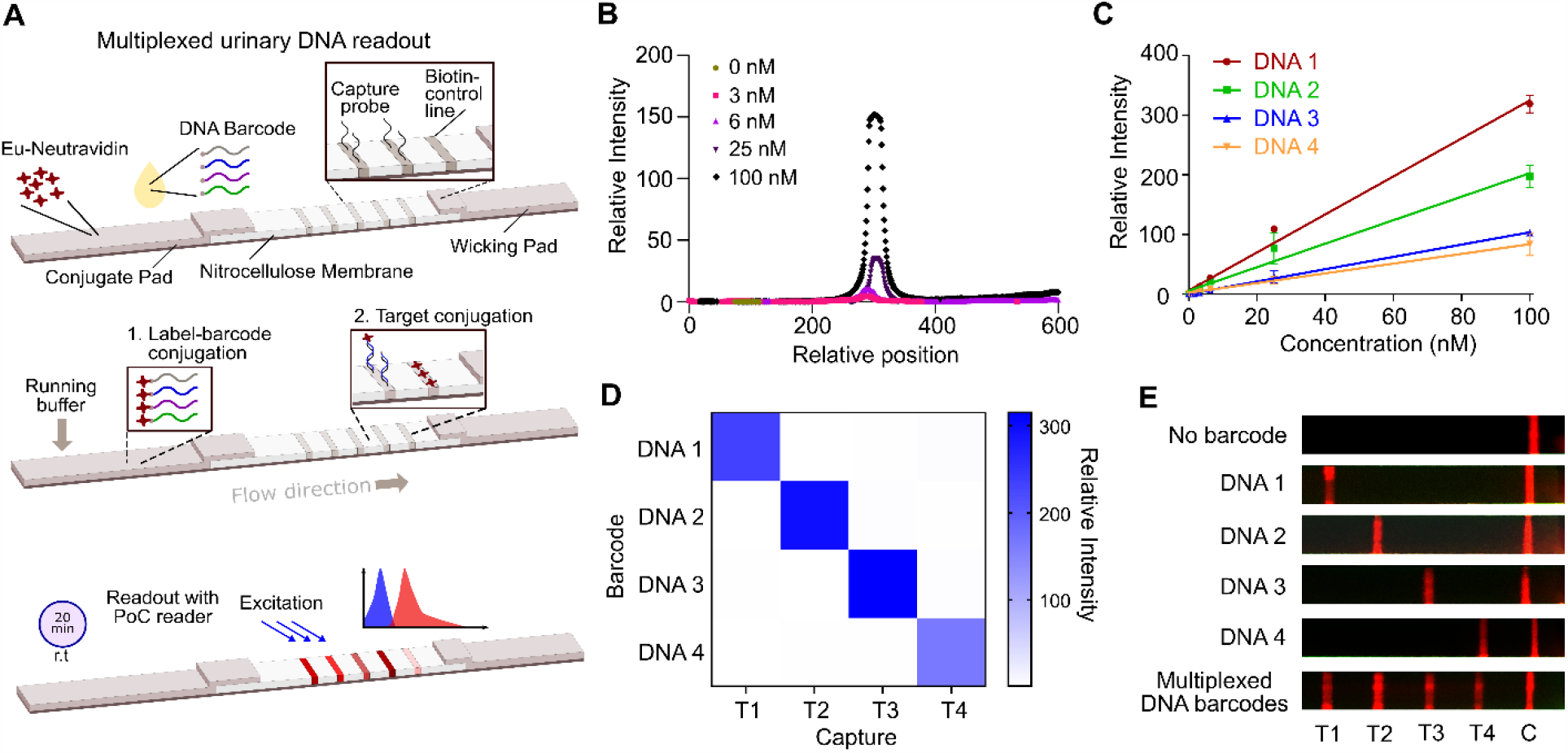
Lateral flow assay (LFA) for the direct multiplexed detection of urinary DNA barcodes at room temperature. **(A)** (top) Orientation of the LFA, showing the conjugation pad, where urine sample and running buffer are applied, the nitrocellulose membrane, where complementary-coded DNA capture probes are immobilized in detection lines, as well as a line of immobilized biotin is included as a control line, the wicking pad which acts as a sink for the reagents. To operate the LFA, urine samples are first spotted onto the conjugate pad with at least a 5mm distance away from the spotted Eu-Neutravidin (middle). With the application of running buffer, Eu-Neutravidin are conjugated with the biotinylated reporter fragments on the conjugate pad and flow laterally to the right, where labelled DNA fragments are captured via sequence-specific hybridization (bottom). After 20 minutes, the FLA strip is inserted into a point-of-care reader, which applies fluorescent excitation and detects the relative intensity of the resulting wavelengths at each designated line location. **(B)** Fluorescence scans across the lateral flow assay after the application of varying concentrations of a specific DNA barcode to a strip with a single detection line. The relative intensity was calculated by subtracting the baseline fluorescent signal. (**C)** Operating linear range of four test lines showing immobilized capture probe for each DNA sequence. All barcodes show a linear response from 0 – 100 nM. Relative intensity of each DNA barcode across the concentration range of interest was normalized against its value at 0 nM. The DNA barcodes and peptide substrates can be randomly paired for synthesizing the DNA-barcoded ABNs. **(D)** Selectivity assay of the LFA developed when run against the four different DNA barcodes of 100 nM. Intensity of each test lines were quantified and compared. **(E)** Photograph demonstrating the multiplexing capabilities of the lateral flow assay developed (under UV lamps). Individual test strips were all fabricated with 4 test lines (T1, T2, T3, and T4) and a control (C), biotin line. Six representative test strips are shown, where the sample administered is indicated to the left of the strip.

We optimized the physical components of the LFA to optimize the sensitivity of the measurement. Specifically, we found that the choice of surfactants used in the running buffer could result in a detrimental effect on the noise level observed on nitrocellulose membranes. As part of the signal optimization process, we screened 24 surfactants against membranes of different pore size and chemical treatment, including commercially-available FF80HP, FF80HP Plus and CN95 (**Fig. S6A**). We first nominated FF80HP Plus with tween 60 as the surfactant in the running buffer since it resulted in low background fluorescence and minimal conjugate aggregation on the strip (**Fig. S6A, B**). We then studied various commercially-available conjugate pads made of microglass, chopped glass and polyesters to contain the Eu-labeled neutravidin. The conjugate pads were laminated onto the nitrocellulose membrane and tested for the release of the Eu conjugates (**Fig. S7A**). Through further optimization, we found that by switching to FF120HP Plus -which has a slower flow rate than FF80HP Plus -the fluorescent signal-to-noise ratio of the oligonucleotide barcode was enhanced on the assay. This improvement could be due to the prolonged interaction time between the barcode and the immobilized capture probes on the membrane. (**Fig. S7B**). **Fig. 5B** shows the relative intensity of fluorescent signals across positions along the nitrocellulose membrane on which a capture probe was hybridized in a single line, after applying varying concentrations of the corresponding DNA barcode. Notably, the assay was sufficiently sensitive to quantify sub nM concentrations of the target. We blocked the Eu-neutravidin with casein and further adjusted the pH of the running buffer to improve signal performance. We were also able to demonstrate the uniform release of the conjugates into the nitrocellulose membrane with minimum aggregation and background (**Fig. S7C**). In summary, the LFA optimized for the quantitative multiplexing of ssDNA barcodes operates with FF120HP Plus as the nitrocellulose membrane, STD17 as the conjugate pad and 0.5% Pluronic F68 in the running buffer of pH 7.4.

The multiplexing capability of the assay was achieved through the spatial printing of individual capture DNA lines, 4 mm apart. We redesigned the dimensions of the LFA (**Fig. S8A**) to be compatible with the commercially-available POC fluorescent reader and benchmarked the performance of the reader against that of a benchtop model in the concentration ranges of interest (**Fig. S8B**). We assayed the detection of varying concentrations of the 4 individual pre-designed ssDNA barcodes (DNA1, 2, 3 and 4 in **Table.S3**) when applied to the 4-plex capture strips, and found that the multiplexed LFA had an operating linear range from 0 -100 nM of DNA1, 2, 3, and 4 (**Fig. 5C**). It is noted that the sensitivity of the assay for each barcode varies within the linear range, which could be due to 1) the hybridization efficiencies of the DNA targets to their respective captures and 2) the amount of capture probes on each test lines. We then tested the selectivity of the LFA. We challenged the assay with solutions containing 100 nM of each individual DNA barcode and analyzed the relative fluorescence using the POC fluorescent reader between each test. As shown in **Fig. 5D**, the assay demonstrated high selectivity of the target DNA barcodes. **Fig. 5E** shows the representative images of the LFA when tested against urine samples spiked with individual or pooled DNA barcodes captured with a smartphone under an UV lamp. In summary, we demonstrated that the customized LFA is able to simultaneously capture quantifiable amounts of multiplexed DNA barcodes in urine with high sensitivity and specificity.

### PATROL integrates two POC technologies for early detection of LUAD

With the individual pieces validated, we sought to integrate the three modules into a single low-resource capable diagnostic platform -PATROL (**Fig. 1, Fig. 6A**). The DNA-coded ABNs were synthesized by coupling biotinylated DNA barcodes to PEG scaffolds via the peptide substrates (i.e. PLQ38-DNA1, PLQ46-DNA2, PLQ81-DNA3 and PLQ83-DNA4). The ABNs were approximately 15 nm in diameter and were highly negatively charged (refer to **Fig. S9** for characterizations). Each PEG nanoscaffold was covalently bonded with approximately 4 tandem peptide-DNA barcodes (**Fig. S9D**). We then continued to utilize the KP LUAD mouse model (KP_7.5wk_) and attempted to validate urinary DNA reporter detection via the LFA after administering nebulizer-delivered ABNs (**Fig. 6B**). **Fig. 6C** shows representative photographs of the LFA for a healthy (blue, top) or KP (red, bottom) mouse, as well as the quantification of the fluorescence signals (measured as area under the curve of each peak) for each of the DNA barcodes. With the peak areas, we calculated the concentration of each barcode in the urine using established standard curves (**Fig. 5C**).

**Figure 6.**
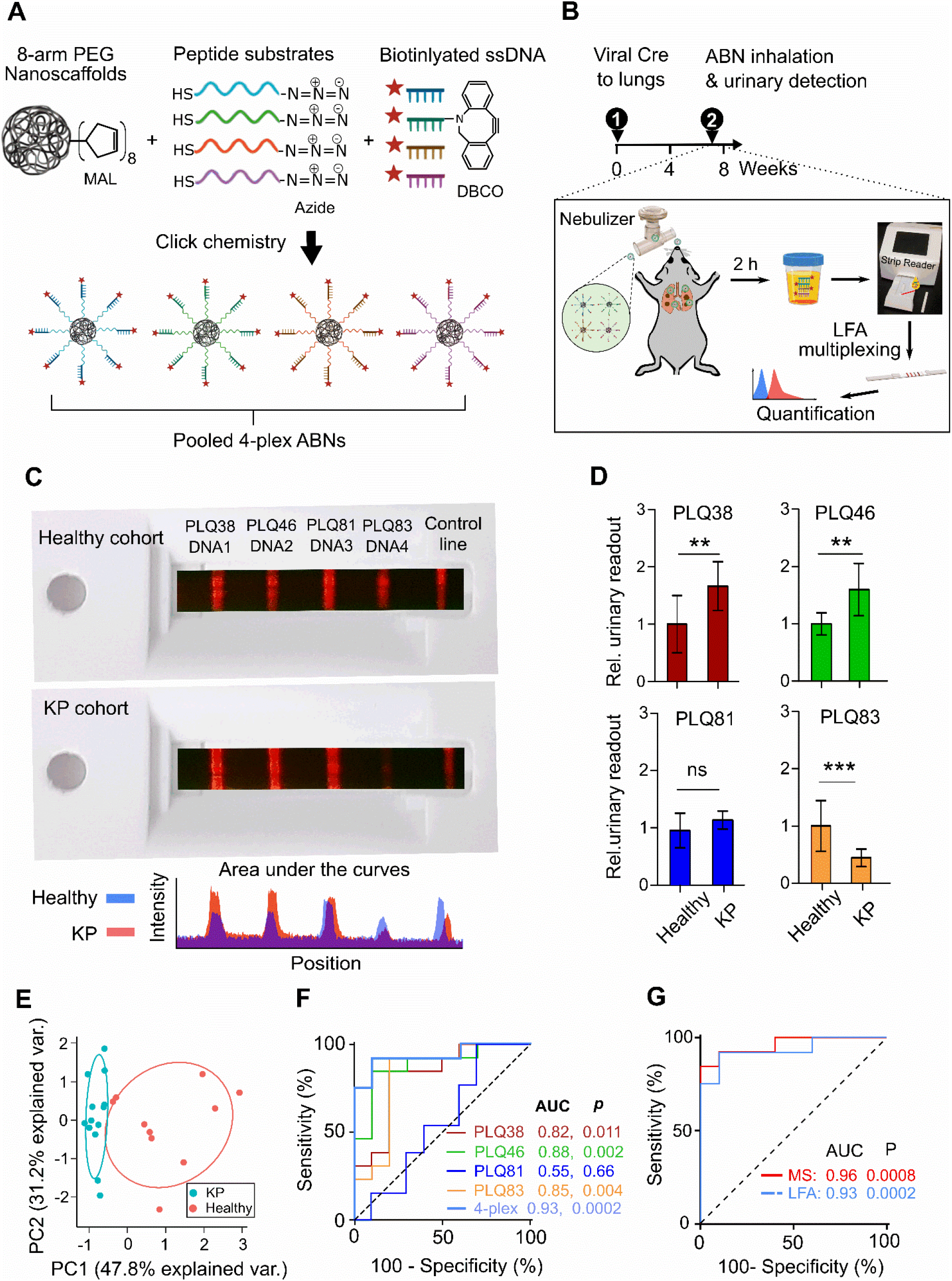
PATROL enabled sensitive classification of LUAD at early stage. **(A)** Single-stranded, phosphorothiolate-modified DNA 20mers and selected protease substrate linkers were conjugated to 8-arm PEG nanoscaffolds via click chemistry, creating a cohort of 4-plexed ABNs with synthetic DNA barcodes. **(B)** Scheme of the workflow for the nebulization of 4-plexed, DNA-encoded nanosensors into the KP_7.5wk_ LUAD and healthy animals, followed by direct detection of synthetic DNA reporters in urine samples using the customized lateral flow assay. **(C)** Photographs demonstrating the detected urinary DNA reporters from KP_7.5wk_ and healthy mice with the customized lateral flow assay. The images of the lateral flow assays (taken under UV lamp) are overlayed on cassettes (under normal lighting conditions) for the point-of-care reader as a proof-of-concept demonstration. Intensity profile across the representative strips was measured with ImageJ. **(D)** Urinary output of 4-plexed DNA-coded nebulized ABNs (PATROL) administered to KP and healthy mice at 7.5 weeks after tumor induction, and detected via LFA fluorescence (KP: n = 13; healthy control: n = 10); *p < 0.05, **p < 0.01 and ***p<0.001 by Student’s t test. Error bars represent SDs. **(E)** PCA of mean-normalized urinary DNA reporters for KP_7.5wk_ mice and healthy controls (KP: n = 13; healthy control: n = 10) administered and detected via the PATROL platform. **(F)** ROC curves showing a predictive power of a single or multiplexed urinary reporters from a subset of KP7.5wk and healthy controls in discriminating an independent test cohort of KP lung tumor from healthy controls. P value represents the significance level between the AUC of KP ROC curves and that of baseline ROC curves. **(G)** AUC of ROC curves with urinary DNA reporters detected using LFA was comparable to that detected with advanced LC-MS/MS.

For the cohorts of mice tested, we compared urinary concentration of each barcode between the 2 groups and plotted the ratios of urinary readouts from KP relative to healthy mice (**Fig. 6D**). We observed significant differences in the cleavage of PLQ38, PLQ46, and PLQ83 but not in PLQ81 between the healthy and KP mice. Unsupervised clustering was able to classify all the KP_7.5wk_ mice with Grade I/II LUAD (**Fig. 6E**). We again performed the ROC analysis to characterize the predictive power of each probe and their combinations. PLQ38, PLQ46 and PLQ83 showed competent predictive power as a single classifier, based on calculated AUC values of 0.82, 0.88 and 0.85, respectively (**Fig. 6F**). The overall AUC of the 4 combined probes increased to 0.93, which was comparable to that of the nebulized 4-plexed ABNs detected via their mass barcodes (AUC=0.96), although the significance (p value) of each probe (except PLQ83) was one order of magnitude lower in the LFA outputs (for DNA) than in the urinary readouts by mass spectrometry (**Fig. 6G**). With 100% specificity, the sensitivity of the DNA reporters detected by the LFA reached 75.2%, slightly less than that of urinary mass reporters, but remains comparable to the sensitivity (i.e. 75%) of microCT that detected LUAD in 7.5 week KP mice at the same specificity.^23^ In summary, diagnostic capacity is retained after switching to PATROL.

As an additional pre-clinical step, we examined the safety profile of a high dose of inhalable ABNs with DNA barcodes (3-fold higher doses than tested above). Neither general toxicity nor clogging of vasculature was observed in major organs of wild-type mice 7 days after a single dose of ABNs delivered via nebulization, as indicated in weight monitoring (**Fig. S10A**) and histological assessment of principal tissues (**Fig. S10B-G**). We further evaluated IFN-γ secretion, a major inflammatory cytokine produced by activated immune cells, and considered an indicator of immunogenicity of an administered substance. At both shorter (i.e 7 days) and longer (14 and 30 days) intervals after ABN dosing, peripheral blood mononuclear cells were isolated and an ELISPOT assay was performed. Upon stimulation with the same panel of the ABNs, we found no IFN-γ was secreted by circulating lymphocytes at any tested time points, consistent with the interpretation that each module of the inhaled ABNs (i.e., peptide substrates, PEG scaffolds, and DNA barcodes) was non-immunogenic (**Fig. S11A**,**B**).

## Discussion

Although there has been substantial progress in lung cancer diagnosis and treatment over the past several decades, progress has not kept pace in tackling cancer health disparities, including low rates of screening and early detection in low socioeconomic groups and geographically isolated areas. A chief solution to address such inequity is to lower the technological threshold for a patient’s access to screening programs and early detection. This work established PATROL as a POC detection platform that integrates modular inhalable synthetic biomarkers and multiplexable LFAs to detect, in a non-invasive manner with low-infrastructure needs, dysregulated proteolytic activity that correlates with early-stage lung cancer (**Fig. 1, Fig. 3B**,**E**). We first formulated the ABNs into microscale aerosols that could be delivered with clinical nebulizers or inhalers and are conducive to deep lung deposition. We demonstrated the inhaled ABNs maintained robust in vivo diagnostic power in a genetically engineered mouse model of lung cancer. Next, we downsized a large library of stage-tailored ABNs to a 4-plex cohort and re-engineered them with synthetic DNA barcodes to enable easy multiplexing with LFAs at room temperature. Finally, we validated that the highly modular ‘inhale and detect’ PATROL platform discriminated Grade I/II LUAD in an autochthonous model with high sensitivity and specificity (AUC=0.93, **Fig. 6F,G**). Collectively, PATROL holds great clinical potential not only to attain both sensitive and specific lung cancer detection at early stages, but also to enable easy deployment in resource-limited settings as demands surge.

Activity-based synthetic biomarkers that selectively leverage protease activity in tumors have yielded amplified signals and outperformed blood biomarkers in detecting smaller tumors in several preclinical models, such as carcinoembryonic antigen (CEA) in colorectal cancer,^32^ and human epididymis protein 4 (HE4) in ovarian cancer.^33^ In addition to output augmentation, PATROL increased detection specificity of the protease-responsive probes by identifying protease signatures that stratify early-stage LUAD using TCGA transcriptomic data (**Fig. 3B, Fig. S3C**). Finally, *in vivo* screening of nominated ABNs results in a robust low-plex probe set for POC assay compatibility. We envision that recent advances in precision multi-omic sequencing will shed new insight into unique protease dysregulation and heterogeneity in various stages and subtypes of lung cancer. The delivery of ABNs via inhalation provides a means of maximizing their deposition to the lungs and decreases background signals caused by non-specific enzymatic activation in circulation. The fully non-invasive inhalation delivery also sets the platform a step closer to clinical translation for direct delivery of diagnostic agents to the lungs. The relatively homogeneous deposition of inhaled ABNs to the entire respiratory system (**Fig. 4A, Fig. S4B**,**C**) may also confer the capacity to profile protease activity of tumor nodules at unknown locations or within the otherwise-inaccessible periphery of the lung. By tailoring aerodynamic size of formulations and rational selection of inhalational devices for subjects, it should be feasible to push the particle deposition down the tracheobronchial tree to alveolar regions, or reverse the deposition pattern to favor the upper airways (**Fig. 2E,F,J,K,L**).

Conventional LFAs often face limitations in quantitative high-throughput analysis of biomarkers. To overcome this shortcoming, PATROL utilizes the most relevant protease substrates to maintain the specificity of a low-plex assay and leverages 20-mer ssDNA sequences as barcodes, which could be also amended for higher-order quantitative multiplexing. Dedicated designs of DNA sequences and the optimization of modules into LFA ‘kits’ (**Fig. S8**) enabled us to detect and quantify urinary barcodes at nanomolar levels (**Fig. 5C,D**). Sequences of 20 nucleotides offer the capacity to extend the platform to hundreds of orthogonal codes, and thus enable us to quickly ramp up multiplexing capacity when needed.^31^ The synthetic DNA barcodes excreted into the urine are quickly captured on paper strips via direct hybridization with their complementary sequences at room temperature. Hence, the LFA tests offer convenient operation without the need for amplification steps used in polymerase chain reactions and CRISPR-based methods, and can be completed within 20 minutes in ambient conditions.^34,35^ Compared to the measurement of mass barcodes by LC-MS/MS, we observed a slight sensitivity decrease in terms of the fold change of some urinary reporters observed between tumor-bearing and healthy mice, such as PLQ81 (**Fig. 4C**,**6D**). Given that the hybridization efficiency of captured DNA varies within a linear range (**Fig. 5B, C**), we envision that pairing LQ81 with more sensitive DNA barcodes (e.g. DNA1) may help amplify the relatively nuanced differences in the activity of some target proteases between tumor and healthy groups. Moreover, the optimization of DNA reporters and capture sequences via an iterative processes of sequence design and validation at *in vitro, ex vivo* and *in vivo* levels would further enhance their binding affinity. In contrast to conventional pregnancy tests and COVID-19 rapid antigen tests where a positive or negative readout is a quantum yes/no result, our multiplexed LFAs require the relative quantification of multiple barcodes with which unsupervised algorithmic methods are used to differentiate a cancer diagnosis from healthy groups. Therefore, for future clinical applications, it is essential to define reliable baselines of these urinary DNA barcodes in lung cancer-free groups, often in the context of other benign or non-cancerous comorbidities.

While these results highlight the encouraging potential to utilize PATROL for screening and early detection of lung cancer, future work remains to be accomplished to improve PATROL’s performance. First, the inhaled ABNs mainly profile protease activity at the periphery of tumor nodules. By enhancing the capacity for PATROL to penetrate into deep tumor locations, we may not only increase the magnitude of detected signals, but also enable the sensors to report unique proteolytic signatures present in the tumor cores. In addition, ABNs can be fine-tuned so as to optimize surface presentation of their substrates^33^ in combination with active targeting to tumor cells and/or microenvironments may further enhance detection sensitivity to monitor for minimal residual disease or early-stage lung cancer. Additional engineering efforts on peptide substrates may further improve the specificity to LUAD. For instance, a tandem substrate with an AND logic gate could enable the liberation of DNA barcodes only upon dual cleavage by two concurrently dysregulated TME proteases. Moreover, the genetically-engineered mouse models utilized in this proof-of-concept report represent only a histological subset of human lung cancer (i.e. adenocarcinoma). Additional efforts must be undertaken to test other subtypes, such as squamous or large cell carcinomas, and to further establish the utility of PATROL for detecting more heterogeneous forms of human lung cancer, also often accompanied by chronic comorbidities. Secondly, despite the finding that DNA barcodes were cleared from murine lungs within 48 hours of administration via nebulizer (**Fig. S11C**), longitudinal safety studies of the PEG, synthetic peptides, and DNA reporters that make up PATROL nanoparticles should receive comprehensive scrutiny.^23^ Another modification that might help reduce the regulatory threshold is to utilize endogenous materials to construct the nanoscaffold (e.g. phospholipids). Finally, although the current PATROL LFAs measure synthetic DNA barcodes in the urine with low-level multiplexing on a single strip, further product development may focus on the expansion of multiplexing capacity and consider clinical uses in decentralized settings. For example, the design of custom-made cassettes could take advantage of the advancement of printing and manufacturing capabilities to expand multiplexing capacity on the same device. Adaptors that offer connectivity to smartphones or wearable devices that enable fluorescence readouts may also be developed to allow for even more robust POC use.^36^

In summary, highly modular diagnostic kits such as PATROL that can offer test results in a single session would be transformative in the detection of lung malignancies. PATROL may also enable efforts to improve risk stratification of early lesions to determine the clinical selection of follow-up procedures. The versatility and modularity of PATROL could also be applied to other pulmonary diseases where proteolytic activities are dysregulated or diagnosis is based on exclusion, such as idiopathic pulmonary arterial hypertension. We envision that by releasing disease screening from its current resource-intensive environment, we may enable feasible surveillance testing that would identify a disease when it is still easy to treat.

