## Supplemental Information for "Inhalable point-of-care urinary diagnostic platform"

[1.] Koch Institute of Integrative Cancer Research, Massachusetts Institute of Technology, Cambridge, MA 02139.
[2.] Marble Center of Cancer Nanomedicine, Massachusetts Institute of Technology, Cambridge, MA 02139.
[3.] Harvard-MIT Division Health Sciences and Technology, Cambridge, MA 02139.
[4.] Institute of Medical Engineering and Science, Massachusetts Institute of Technology, Cambridge, MA 02139.
[5.] Department of Biology, Massachusetts Institute of Technology, Cambridge, MA 02139.
[6.] Howard Hughes Medical Institute, Massachusetts Institute of Technology, Cambridge, MA 02139.
[*] These authors contributed equally to this work.

### Materials

All chemical reagents were purchased from Millipore-Sigma (Burlington, MA) unless specified.

### Methods

**Particle size measurement of nebulized nanosensors by dispersion laser diffraction**

Volumetric size of particles was measured using a Sympatec HELOS/RODOS (Sympatec GmbH, Clausthal-Zellerfel, Germany) employing the rotary feeder and R3 lens (0.9–175 μm). Powder was hand-filled into the u-shaped groove of the rotating table to cover a length of approximately 1 cm. The sample passed under a plough scraper and roller to remove any excess and was subsequently drawn up into the dispersing line via the protruding aspiration tube from a static bed. During sample delivery the rotating table was maintained at a constant rotation setting of 20%. The measurement was set to trigger when the optical concentration (Copt) exceeded 1.1% and cease when the Copt fell below 1% for 5 s (or 60 s real time). The timebase was 100 ms and a forced stability of '4' was applied. The primary pressure (PP) was manually set using the adjustment valve in the range 0.2–4.5 Bar and three measurements were taken at each pressure setting using freshly loaded powder. PSDs (Dv10, Dv50, Dv90 and VMD) were calculated using Fraunhofer theory and analysed in WINDOX 4.0 software. Particle size measurements for a complete titration curve were made on a single day.

**Particle sizing measurement and fine particle dose of nebulized ABNs using next generation impactor**

The aerodynamic size of inhalable ABN formulations was determined with a next generation impactor (Copley Scientific, Nottingham, UK). The pre-separator and plates were precoated with silicone greases to minimize the bounce of particles on their surface. Mass-coded, Cy7-labled model ABN ($PEG_8$40k-LQ81-GluFib) in 0.5 mL normal saline at different concentrations were loaded into an Aeroneb vibrating-mesh nebulizer at different concentrations ($PEG_8$40k concentration: 25, 62.5 and 200 μM). The solution was nebulized into next generation impactor (NGI) at a flow rate of 15 L/min (Fig.S1B, C). The nebulizer (bottom part) and T tubing, induction port (IP), pre-separator (PS), 8 stages, and nylon filter were carefully rinsed with 3 mL deionized $H_2O$, and then measured for Cy7 fluorescence at Ex/Em = 735/780 nm (note: a necessary dilution may be required for some samples). Fine particle fraction (FPF), the fraction of particles with aerodynamic diameter<5.0μm implying the fraction of aerosols that tend to deposit on peripheral lung regions (Fig.2D), was calculated with the following equation:

$$\mathrm{FPF} = \frac{\text{Dose of ABNs with particle size} < 5\ \mu\text{m}}{\text{Total dose of drug deposited on IP, PS, stage } 0-7, \text{and MOC}}$$

Mean mass aerodynamic diameter (MMAD), the aerodynamic particle size of the cumulative percentage of 50% in cumulative particle distribution, was obtained by plotting probability (cumulative percentage of deposited aerosols on stages) against log (upper cutoff diameter of each).

**Spray drying of ABN-laden Microparticles.** ABNs (4mg), D-mannitol (12 mg) and L-leucine (4 mg) were dissolved in 5 mL deionized (DI) $H_2O$ and then spray dried using B-290 Mini Spray Dryer (BUCHI) with the following parameters: inlet temperature = 60 C, outlet temperature = 39–44 C, atomizing nitrogen flow rate = 35 mm (473 L/h), pump ratio = 10% (3 mL/min), aspiration rate = 70% (100% = 35 m3/h), and nozzle cleaning 0. The spray-dried microparticles were collected at the end of the gas cyclone. The resulting microparticles were further dried overnight to remove residual $H_2O$ in a FreeZone benchtop freeze dryer (Labconco) with the cold well set at −80 C and the vacuum degree at 0.02–0.015 mbar. The dried samples were then stored in a Corning® PYREX® top glass desiccator sealed with silicone lubricant, and Drierite® was used as a desiccant.
For dry powder inhalers,

$$\text{Emitted dose} = \frac{\text{Dose of ABNs released into NGI}}{\text{Total dose of ABNs loaded into a capsule}}$$

, and FPFs and MMAD were calculated as mentioned above.

**Nanosensor stability and activity post nebulization**

Analysis of nanoparticle stability following nebulization used 0.1 mL 200 μM Cy7-labeled LQ81 $PEG_8$40kDa nanosensors (Fig.2G, H). The ABN was nebulized into a thoroughly washed glass beaker. The collected nanosensors were measured for hydrodynamic size and zeta potential with a NanoZS Zetasizer (Malvern). Protease cleavage assays of aerosolized nanoparticles used LQ81 $PEG_8$40kDa nanosensors with FAM and QSY21 as a quenched FRET pair. The nebulized nanosensors were diluted to 1 μM and incubated with 12.5 nM recombinant human MMP13 (Enzo). Proteolytic cleavage of substrates was quantified by increases in fluorescence over time by a Tecan Infinite M200 Pro fluorimeter. Enzyme cleavage rates were quantified as relative fluorescence increase over time normalized to fluorescence before addition of protease, and the results were benchmarked against untreated nanosensors.

**Oral gavage of ABNs**

Cy7-labeled LQ81-GluFib peptide with a cysteine at C-terminus was synthesized by CFC scientific. The peptide was then conjugated to a 40kDa 8-arm PEG maleimide nanoscaffold (Jenkem Technology) overnight at room temperature as a model ABN. The resulting crude product was then first concentrated to approximately 0.5 mL with an Amicon 0.5 mL, MWCO 10kDa, ultra-centrifugal filter (EMD Millipore), followed by the purification using a Cytiva ÄKTA fast-protein liquid chromatography system equipped with a Sephadex 30 Increase 10/300GL column (Groton, CT). The purified model ABN was reconstituted in the PBS (1X, pH 7.4), and diluted to 20 μM (substrate equivalent). A 50 uL of the model ABN in the PBS was administered with a flexible plastic tubing oral gavage needle from GavageNeedle (Phoenix, AZ). One cohort of mice (n=3) was used to collect blood samples retro-orbitally with heparinized capillary tubes at different timepoints. The 40 μL (2-3 drops) blood from each mouse was mixed with 60 μL heparin solution (10U/mL), and the mixture was centrifuged at 1000g. 50 μL plasma supernatant was collected and measured for Cy7 fluorescence with a LI-COR Odyssey® DLx imager (Lincoln, NE). The same dose of ABN was also injected intravenously, and blood was collected as mentioned above for comparison. In parallel, orally gavaged mice were euthanized at 1, 4 and 24 hours, and lung, heart, stomach, small intestine, liver, spleen, kidney and esophagus were carefully excised for the biodistribution evaluation using a LI-COR Odyssey® DLx imager (LI-COR Biosciences).

**Generation autochthonous lung adenocarcinoma on mouse models.**

To induce lung tumors that resemble the human NSCLCs histopathologically, we first generated the K-$ras^{LSL\text{-}G12D/+}$;$p53^{fl/fl}$ C57BL/6 (KP) mice in which the activation of an oncogenic allele of K-ras is sufficient to initiate the tumorigenesis process, and additional deletion or point mutation of p53 significantly enhances tumor progression, leading to a more rapid development of adenocarcinomas that have features of a more advanced disease. Lung tumors were initiated by intratracheal administration of 50 μL containing adenovirus-SPC-Cre ($2.5\text{x}10^8$ plaque-forming units (PFU) in Opti-MEM with 10 mM calcium chloride

($CaCl_2$)) in female or male KP mice (between 8 and 16 weeks old) under isoflurane anesthesia. Control cohorts consisted of age and sex-matched mice that also underwent intratracheal administration of AdenoCre. The KP mice were allowed for tumor growth until the time experiments were performed. All animal studies and procedures were approved by the Committee for Animal Care at Massachusetts Institute of Technology.

**Gene expression analysis**

Human RNA-Seq data from The Cancer Genome Atlas (TCGA) Research Network (32) was downloaded from https://www.cancer.gov/tcga. The list of human extracellular protease genes was obtained from UniProt using the following query: (keyword:"Protease [KW- 0645]") locations:(location:secreted) AND reviewed:yes AND organism:"Homo sapiens (Human) [9606]"). Differential expression analysis on the TCGA data was performed using the DESeq2 differential expression library in the R statistical environment. Differentially expressed genes were filtered to have p-adjusted value <0.05 and to contain only extracellular proteases and were ranked as a function of $\log_2$(fold-change Stage I LUAD/NAT expression) (Fig. 2A). Proteases were color-coded as a function of protease class.

***In vitro* screening of substrate peptides against LUAD-associated proteases at early stages**

Fluorogenic protease substrates were synthesized using Millipose Sigma's custome peptide libray service PepScreen (LQx) or by CPC (PPx, GzmB, NE, CatK) (Table S1, Fig.S2B,C, Fig.3B). Recombinant proteases were purchased from Enzo Life Sciences, R&D Systems, and Haematologic Technologies. For recombinant protease assays, fluorogenic substrates LQx (60 µM final concentration) or PPx, GzmB, NE, CatK (1 uM final concentration) were incubated in 30 µL final volume in appropriate enzyme buffer, according to manufacturer specifications, with 12.5 nM recombinant enzyme at 37°C (Fig.S2B). Proteolytic cleavage of substrates was quantified by increases in fluorescence over time by a fluorimeter (Tecan Infinite M200 Pro). Enzyme cleavage rates were quantified as relative fluorescence increase over time normalized to fluorescence before addition of protease. Hierarchical clustering was performed in GENE-E (https://software.broadinstitute.org/GENE-E/, Broad Institute), using z-scored fluorescence fold changes at 60 minutes.

**Synthesis and characterization of DNA-coded, multiplexed activity-based nanosensors (ABNs)**

All aqueous buffers were deoxygenated by a succession of 30 min vacuum and 15 min dry nitrogen purge. Each of azide-functionalized peptide with a C-terminal cysteine (CPC Scientific, San Jose, CA) was dissolved in dimethylformamide (DMF) to reach 10 mg/mL. A 20 µL of the peptide solution was then slowly added to 380 µL phosphate buffer (pH 7.0, 0.2 M), followed by an addition of 500 uL of 200 µM DNA barcode (Integrated DNA Technologies, Coralville, IA). The solution was stirred slowly under the protection of dry nitrogen at room temperature for 1 hour. A 100 µL phosphate buffer containing 0.82 mg 40 kDa 8-arm PEG maleimide (Jenkem Technology, Beijing, China) were then added dropwise and the reaction was stirred overnight at room temperature. The resulting crude product was then first concentrated to approximately 0.5 mL with an Amicon 0.5 mL, MWCO 10kDa, ultracentrifugal filter (EMD Millipore, Burlington, MA), followed by the purification using a ÄKTA fast-protein liquid chromatography system equipped with a Sephadex 30 Increase 10/300GL column (Cytiva, Groton, CT). The fractionates eluted from 6.5 mL to 10 mL were combined and concentrated with an Amicon 0.5 mL ultracentrifugal filter (MWCO 10KDa). The molar concentration of DNA was determined with the Quant-iT™ OliGreen™ ssDNA Reagent kit (Thermo Scientific, Waltham, MA) according to a manufacturer's protocol.

Mass-assisted laser desorption/ionization-time of flight mass spectrometry (MALDI-TOF) was performed on a Bruker UltrafleXtreme mass spectrometer (Bruker Corporation, Billerica, MA) to confirm the molecular weights of necessary intermediates and purified products. Sinapinic acid was used as a matrix compound for MALDI-TOF. Hydrodynamic sizes and surface charges of ABNs were measured using a

Malvern Zetasizer Nano ZS (Malvern, United Kingdom). The geometry and size of ABNs was also visualized with a JEOL2100F cryo-transmission electron microscope (Tokyo, Japan).

**Intratracheal administration of 20 multiplexed ABNs to the KP mice**

A library of 20-plexed, mass-coded ABNs by enriching the peptide reporter glutamate-fibrinopeptide B (GluFib) (EGVNDNEEGFFSAR) with different distributions of stable isotopes for *in vivo* experiments (Table S2) were synthesized and characterized by CPC Scientific (Sunnyvale, CA). All *in vivo* administration of ABNs experiments were performed in the morning and were in strict accordance with institutional animal guidelines. Bladders of mice were voided immediately before nanosensor administration. Mice were anesthetized with isoflurane inhalation (Zoetis, Parsippany-Troy Hills, NJ), and were monitored during recovery. For intratracheal instillation studies, a volume of 50 µL of multiplexed (20-plex or 4-plex), mass-coded or DNA-coded ABN cocktail (20 µM substrate equivalent) in phosphate buffer (0.28 M mannitol, 5 mM sodium phosphate monobasic, 15 mM sodium phosphate dibasic, pH 7.2-7.4) was administered by solution injection following intratracheal intubation with a 22G flexible plastic catheter (Exel International, Redondo Beach, CA) and a 100 µL of air was then pushed into mouse lung lobes via a 1 mL syringe to ensure a full delivery of ABNs into the lungs. A subcutaneous injection of 200 µL sterile PBS was followed in favor of urine production. Bladders were voided again at 60 minutes after nanosensor administration, and urines produced from 60 to 120 minutes post administration were collected using custom tubes in which the animals rested on 96-well plates that collect urines. The urine on every plate was pooled together as a sample from one mouse. All urine samples were immediately frozen at -80°C until analysis by LC-MS or by customized lateral flow assay (LFA) kits.

**Aerosolization of multiplexed activity-based nanosensors (ABNs)**

Exposure of KP and healthy mice to aerosols containing 4-plex, mass-coded or DNA-coded ABNs were performed in the nose-only inhalation tower system (CH Technologies, Westwood, NJ), including an aerosol distribution chamber and a number of animal containment tubes. Mice were first acclimated to containment tubes by keeping them in the tubes for 15 minutes and twice a day for 3 consecutive days before inhalation experiments. Such training mitigates animals' stresses and ensures normal breathing frequencies and cycles during the restraint. In the morning of experiment day, bladders of the mice were voided of urines and then gently loaded into the tubes that were connected to the ports of the inhalation tower. Four-plex ABN cocktails (125 µM per nanosensor) in 0.5 mL normal saline were added into an Aeroneb® mesh-vibrating nebulizer (Aerogen, Chicago, IL) and the ice were exposed for 15 to 20 minutes to aerosols containing ABNs that were generated via the nebulizer and drifted into the tower along with dry airs at a flow rate of 1 L/min. To determine inhaled dose, an all-glass impinger (AGI; catalog no. 7541-10, Ace Glass, Vineland, NJ) containing 10 mL of PBS + 0.001% antifoam was attached to the chamber and operated at 6 lpm, −6 to −15 psi. Particle size was measured once during each exposure at 5 min using the Aerodynamic Particle Sizer (TSI, Shoreview, MN) operating at 5 lpm. A 5-min air wash followed each aerosol, after which animals were returned to their cage. AGI samples were evaluated to determine the concentration of nanosensors recovered from the aerosol. The inhaled dose was determined as the product of the Nb aerosol concentration, duration of exposure, and the minute volume of individual mouse. Minute volume was determined using Guyton's formula. A subcutaneous injection of 200 µL sterile PBS followed aerosol delivery of nanosensors in favor of urine production. Bladders were voided again 60 minutes after the end of nebulization, and urines produced 60-120 min were collected in the custom tubes. The urines were frozen at -80°C until analysis by LC-MS (for mass-coded ABNs) or by customized, paper-based LFA kits (for DNA-coded ABNs).

**Quantification of mass-coded urinary reporters with LC-MS/MS**

LC-MS/MS was performed by Syneos Health (Princeton, NJ) using a Sciex 6500 triple quadrupole instrument. Briefly, urine samples were treated with ultraviolet (UV) irradiation to photocleave the 3-Amino3-(2-nitro phenyl)propionic Acid (ANP) linker and liberate the GluFib reporters from residual peptide fragments. Samples were extracted by solid-phase extraction and analyzed by multiple reaction monitoring by LC-MS/MS to quantify concentration of each GluFib mass variant. Analyte quantities were normalized to a spiked-in internal standard and concentrations were calculated from a standard curve using PAR to the internal standard. Mean normalization was performed on PAR values to account for mouse-to-mouse differences in ABN inhalation efficiency and urine concentration

**Immobilization of capture probes on the multiplexed lateral flow assay**

Capture oligonucleotides C1, C2, C3 and C4 (free of amine byproducts) synthesized by Integrated DNA Technologies (Coralville, IA) were reconstituted in 500 µL of 0.1 M borate buffer of pH 9.3 (DCN, Carlsbad, CA). The amine group was activated for bovine serum albumin (BSA) (Equitech-Bio, Kerrville, TX) conjugation with 5 mL of p-Phenylene diisothiocyanate (PDITC) dissolved in Dimethyl Formamide (DMF) at a concentration of 20 mg/mL. The reaction mixture was kept in the dark at room temperature followed by the addition of 9 mL of n-butanol and 15 mL of nuclease-free water. The samples were mixed well and centrifuged at 15000g for 5 minutes. The yellow layer was discarded and the extraction process was repeated. The samples were then freeze-dried. To conjugate the activated capture oligonucleotide to BSA, 48.5 nmol of C1, 54.2 nmol of C2, 64 nmol of C3 and 48.5 nmol of C4 were added to 0.20 mL, 0.224 mL, 0.264 mL and 0.2 mL of 2mg/mL BSA respectively. The reaction was done overnight at room temperature in the dark and dialyzed against 10 mM phosphate buffer 4 times. The conjugates C1, C2, C3 and C4 were diluted 4, 16, 8 and 32 times respectively with 10 mM acetate buffer at pH 5.0 (DCN, Carlsbad, CA) and printed on the nitrocellulose test strip (FF120HP Plus) with Biodot (Irvine, CA) programmed to stripe at 0.6 µL/cm @ 10 mm/s. The relative position of capture C1, C2, C3 and C4 were 5 mm, 9mm, 13mm and 17 mm from the edge of the membrane. Likewise, biotin-BSA (DCN, Carlsbad, CA) of 0.15 mg/mL were also printed on FF120HP Plus (Cytiva, Malborough, MA) at 21 mm away from the edge of the membrane (control line). The membranes were dried at 40°C for 30 minutes before placing them into a foil pouch with desiccants (DCN, Carlsbad, CA). It is noted that C1, C2, C3, C4 are capture probes for DNA1, DNA2, DNA3, and DNA4, respectively.

**Preparation of the Eu-Neutravidin label**

The solution in the 1% europium (Eu) beads (Thermo Fisher Scientific, Waltham, MA) stock was replaced with 500 µL of 0.1 M 2-(N-morpholino)ethanesulfonic acid (MES) buffer via 3 rounds of centrifugation at 15000g for 15 minutes. 1-ethyl-3-(3-dimethylaminopropyl)carbodiimide (EDC) and N-hydroxysuccinimide (NHS) were dissolved at a concentration of 15 mg/mL and 50 mg/mL in 0.1 M MES (DCN, Carlsbad, CA) at pH 6 before use. 0.1 M of MES at pH 6 was added to the Eu beads solution followed by the addition of 5 µL of EDC and 100 µL of NHS. The reaction mixture was rotated on an orbital shaker with 500 rpm for 30 minutes at room temperature. The mixture was further purified with 3 rounds of centrifugation at 15000 g for 15 minutes. After removing the supernatant, 200 µL of 200 mM phosphate buffer and 250 µL of 1 mg/mL Neutravidin (Thermo Fisher Scientific, Waltham, MA) in 10 mM phosphate buffer at pH 7.2 (DCN, Carlsbad, CA) were added. The solution was mixed on an orbital shaker (500 rpm) for 3 hours at room temperature, followed by the addition of 5 µL of 1 M ethanolamine (TCI America, Portland, OR) and continue to shake overnight. The Eu-Neutravidin label was then resuspended in 50 mM TRIS, 1% casein, 5 mM EDTA, 0.2% Tween-20 of pH 8.0 (DCN, Carlsbad, CA) via 3 rounds of centrifugation at 15000 g for 15 minutes. This allowed the blocking of Eu-Neutravidin label with Casein.

**Spray drying of Eu-Neutravidin on the conjugate pad**

60 µL of Eu-Neutravidin conjugate was added to 1140 µL of 50 mg/mL trehalose in deionized (DI) water. The solution was then printed onto the conjugate pad, STD17 (Cytiva, Malborough, MA), using BioDot Airjet (Irvine, CA). The pads were dried at 40°C for 30 minutes before placing them into a foil pouch with desiccants.

**Lamination of the LFAs**

The membrane was first laminated onto the backing card through adhesive (DCN, Carlsbad, CA). This was followed by the lamination of Ahlstrom 270 (Ahlstrom-Munksjö, Helsinki, Finland) on the top of the card as a wicking pad. The conjugate pads were then placed at the other end of the nitrocellulose membrane and secured with adhesives. The backing card was then trimmed to 5 mm strips to allow the strips to be placed into the cassette of the point-of-care reader.

**Detection and quantification of urinary synthetic DNA barcodes with lateral flow assay**

25 mM TRIS, 150 mM NaCl, 0.5% Pluronic F68 (Thermo Fisher Scientific, Waltham, MA), pH 7.4 was used as the running buffer to perform the measurement. 5 μL of urine samples were first added to the conjugate pad at a position of ~ 5 mm away from the nitrocellulose membrane. A 10 μL of the running buffer was then placed 10 mm below the urine samples as a spacer. The strips were then immersed into 125 μL of the running buffer in a 48 well plate. The strips were left to run for 20 minutes at room temperature to allow the conjugation of DNA barcodes to the Eu-Neutravidin label and also the hybridization at their respective capture probes. The strips were then analyzed with an Axxin AX-2X-S point-of-care reader (Fairfield, VIC, Australia)

**Immunohistochemical staining of ABNs in the lungs**

For immunohistochemical visualization of nanosensors following intratracheal administration and nebulzation, EZLink™ NHS-Biotin (Thermo Fisher Scientific) was coupled to amine-terminated $PEG_8$40k at a 2:1 molar ratio in DMSO in presence of a trace of triethylamine and the reaction was carried out overnight, followed by washes and purification with Millipore Amicon® MWCO 10KDa ultra-0.5 centrifugal filters (Burlington, MA). Pulmonary administration of biotinylated $PEG_8$40k nanoscaffolds was performed by intratracheal instillation (50 μL, 20 μM biotin equivalent) or nebulization (0.2 mL, 50 μM biotin equivalent) as described above. Fixation was performed 2 hours after administration by inflating lungs with 1 mL 4% paraformaldehyde (PFA). Lungs were then excised, fixed in 4% PFA at room temperature overnight, and embedded in paraffin blocks for sectioning. Five μm tissue slices were sectioned and stained for biotin with a streptavidin-HRP ABC kit (Vector Laboratories, Burlingame, CA) following a protocol provided by the manufacturer. The stained slides were scanned using the 20x objective of a 3DHistech Pannoramic 250 Flash III whole slide scanner (Budapest, Hungary), and analyzed with QuPath 0.3.0.

**Examination of general histopathological toxicity**

A set of 4-plex, DNA-coded ABNs were nebulized to wild-type C57BL/6 mice via an inhalation tower system as mentioned above. The body weights of the mice were monitored for 7 days and normalized to that at the day of ABN nebulization. Lung, liver, heart, spleen, kidney and brain were carefully excised, fixed in 4% PFA at room temperature overnight, and embedded in paraffin blocks for sectioning. Five μm tissue slices were sectioned and stained for hematoxylin and eosin in the Hope Babette Tang (1983) Histology Facility at the Koch Institute of MIT. All slides were carefully examined by a highly experienced veterinary pathologist (Dr. Roderick Bronson from Harvard Medical School).

**Immunogenicity of DNA-coded nanosensors by ELISpot**

The IFN-γ enzyme-linked immunospot (ELISpot) assay has been used extensively for the screening of immune responses elicited by foreign antigens. A volume of approximately 0.2 mL blood were sampled into EDTA-coated tubes via retro-orbital bleeding at Day 7, 14 and 30 following the intratracheal administration of 4-plex, DNA-coded nanosensor cocktails (50 μL, 20 μM per nanosensor). Red blood cells were lysed with ACK lysing buffers (Thermo Fisher Scientific) and white blood cells were collected by centrifugation at 300 x g for 5 minutes at 4 °C. The cell pellet was resuspended in cold PBS and washed one more time. The resuspended cells then diluted to 1 million cells/mL with Gibco™ RPMI1640 (Thermo Fisher Scientific) supplemented with 10% fetal bovine serum (Thermo Fisher Scientific), and the cells were seeded on a 96-well plate that was pre-coated with anti-IFN-γ capture antibodies. A pooled 10 μg sterile 4-plex, DNA-coded ABNs were added to the cells and incubated for 24 h. Secreted IFN-γ were detected with a mouse IFN-γ ELISPOT Pair BD Biosciences, Franklin Lakes, NJ) following the manufacturer's protocol.

Mass-coded, Cy7-labled model ABN ($PEG_8$40k-LQ81-GluFib) in 0.5 mL normal saline at different concentrations were loaded into an Aeroneb vibrating-mesh nebulizer at different concentrations ($PEG_8$40k concentration: 25, 62.5 and 200 μM). The solution was nebulized into Helos laser diffraction

### Supplementary Figures

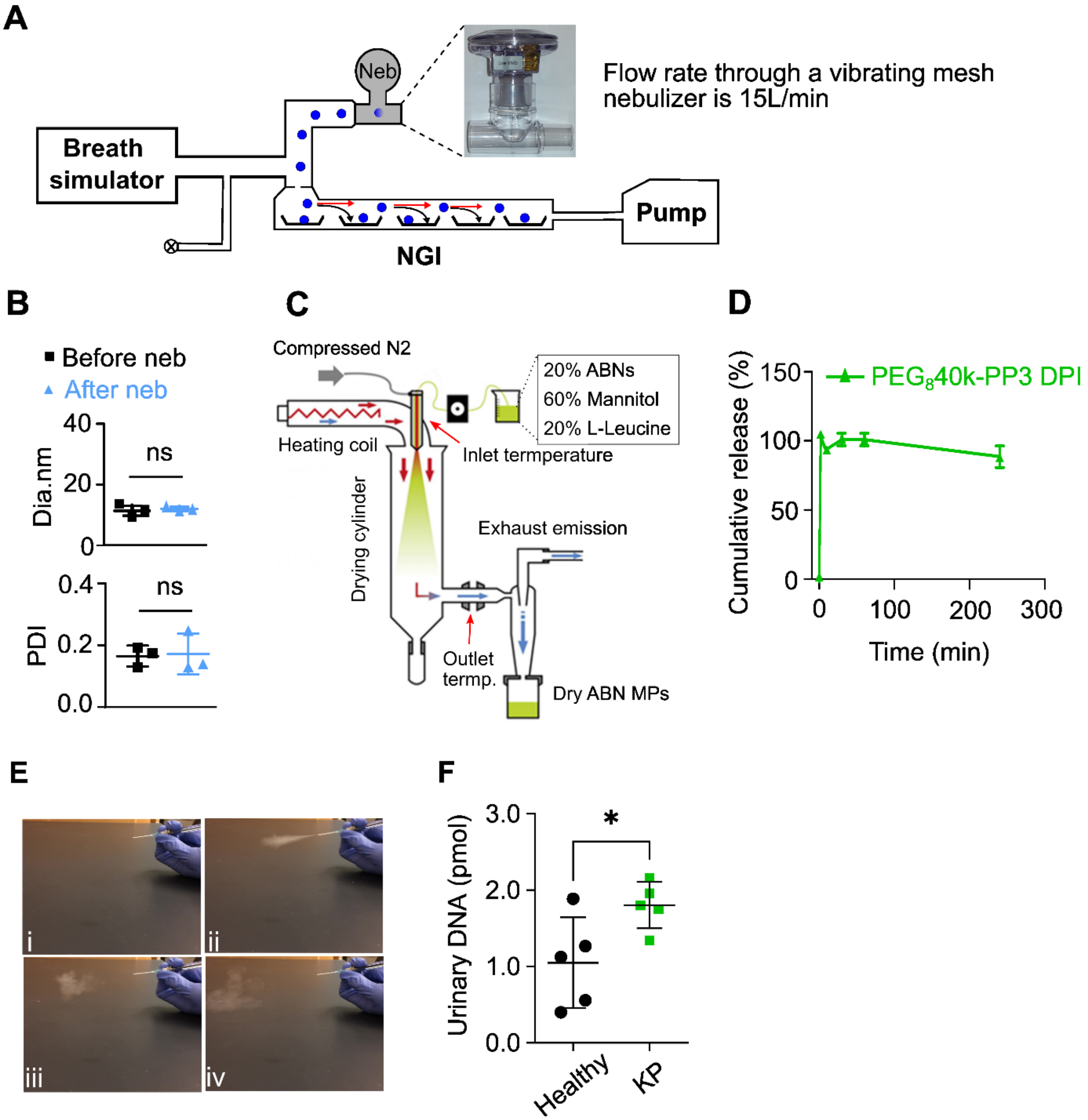


**Figure S1. Characterization of aerodynamic performance of ABN aerosols from vibrating mesh nebulizers. (A)** A breath simulator that generates and applies inhalation-exhalation profiles that mimics that of a human subject was connected to a next generation impactor (NGI) to improve clinical relevance of the nebulized ABNs in in vitro testing. For instance, delivery dose uniformity and fine particle doses. The tests were carried out under a standardized breathing pattern of 500 mL tidal volume, 1:1 inhalation:exhalation (I:E) ratio, and 15 breaths per minute (BPM) frequency. **(B)** Hydrodynamic diameter (top) and polydispersity (bottom) of the model ABN remained basically unchanged before and after vibrating mesh nebulization. Abbreviation: Dia: diameter, PDI: polydispersity index. **(C)** Scheme of spray drying process that is utilized to make ABN-laden microparticles. The ABNs (20% by weight) were sprayed into the microparticles using a Buchi Min-290 spray dryer along with pharmaceutical excipients; D-mannitol (60% by weight) and L-leucine (20% by weight). **(D)** ABNs were immediately released from the microparticles when incubating with synthetic lung fluids. One hundred microliters of solution were added to a 1.5 mL Eppendorf tube and ABNs were isolated by centrifugation. The supernatant was quantified for Cy7 fluorescence. **(E)** The ABN-laden microparticles were aerosolized with a homemade dry powder insufflator**. (F)** Urinary DNA reporters quantified with LFA immobilizing complementary sequence of DNA barcode 3. The ABNs with LQ81-DNA3 were intratracheally delivered with the dry powder insufflator to the mice with or without transplanted tumors in the lungs. The urine samples were collected 2 hours post administration. KP = female C57BL/6 mice with KP tumors by injecting lung epithelial cancer cells isolated from C57BL/6 mice with Kras mutation and p53 inactivation (KP) via tail vein, and tumors were allowed to developed for 2 weeks before dosing the ABNs.

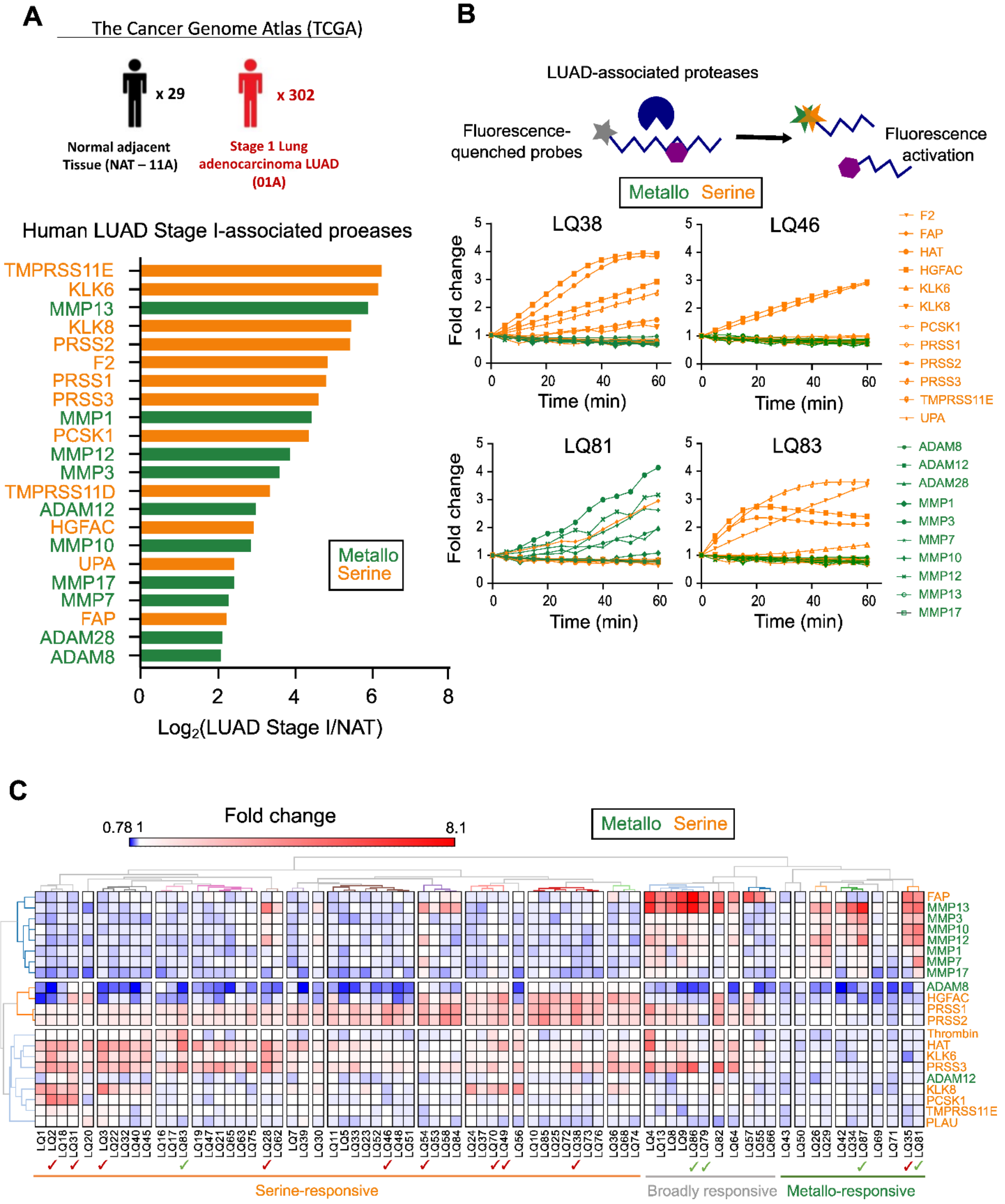


**Figure S2. Overexpressed proteases in Stage I are powerful biomarkers to be leveraged for early detection of lung adenocarcinoma (LUAD). (A)** Differential gene expression analysis between stage I LUAD and non-adjacent tumor tissue (NAT) using The Cancer Genome Data (TCGA) LUAD data identified top 50 dysregulated proteases. Twenty-two proteases of which recombinant counterparts are commercially available were plotted and ranked by fold change. Metalloproteases and serine proteases were color-coded green and orange, respectively. **(B)** A set of 22 proteases dysregulated in Stage I LUAD were screened against of a panel of 15 Förster resonance energy transfer (FRET)–paired protease substrates, and fluorescence activation (green for metallo-proteases and orange for serine proteases) was recorded for 60 min. Fluorescence fold change over 60 min (average of 2 replicates) were z-scored by row and tabulated. Hierarchical clustering was performed to cluster substrates (horizontal) by their protease specificity, Representative kinetic fluorescence curves of LQ38, LQ46, LQ81 and LQ83 against all proteases were shown respectively. The data show that preferential cleavage of LQ81 is by metalloproteases (green) and LQ38, LQ46, LQ83 are by serine proteases (orange). **(C)** Heatmap of z-scored fluorescent fold-changes at 50min (average of 2 replicates) showing hierarchical clustering of proteases dysregulated in Stage I LUAD (vertical) by their substrate specificities and of selected FRET-paired synthetic substrates (horizontal) by their protease specificity. Ticks indicate substrates nominated for in vivo evaluation. TMPRSS11D was excluded from the list due to unavailable recombinant counterpart.

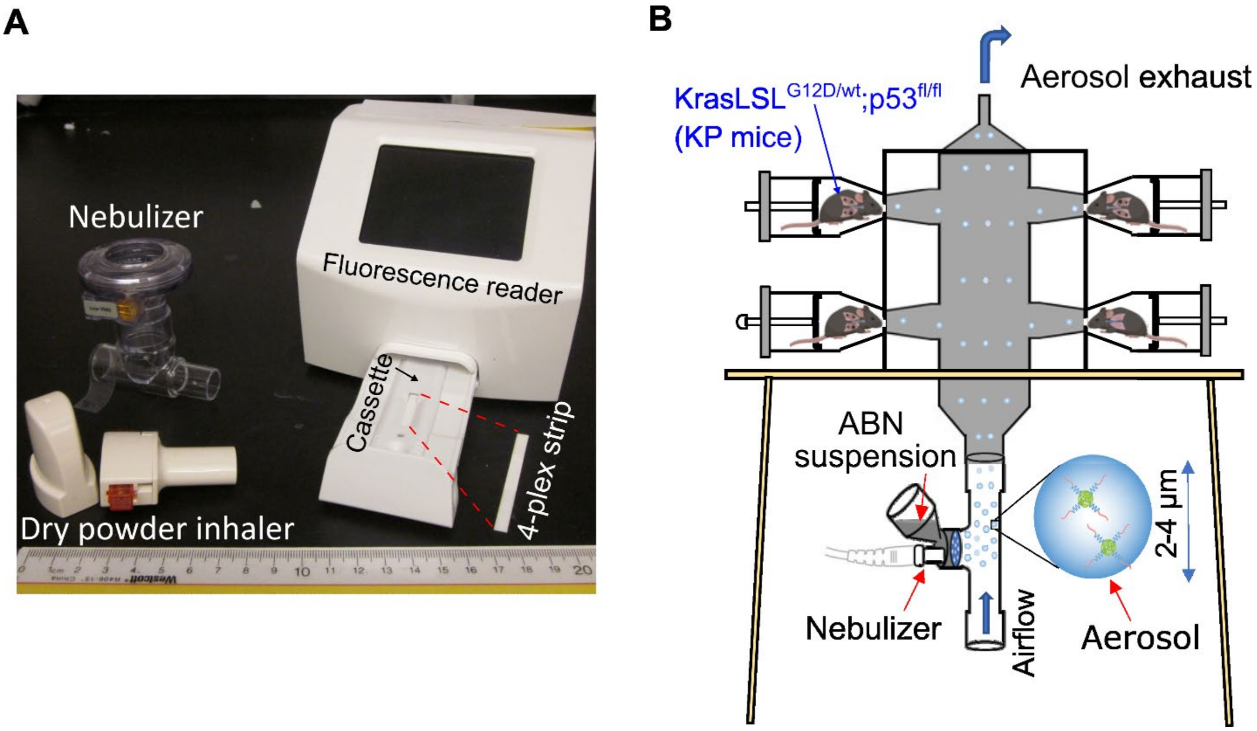


**Figure S3. Prototype components for inhalable point-of-care urinary diagnostic platform. (A**) Inhalable formulations of the ABNs can be generated via either nebulizers or dry powder inhalers for non-invasive pulmonary delivery. Urine samples are collected and then added on a paper strip to multiplex DNA barcode signals quantitatively with a portable lateral flow assay fluorescent reader. Quantitative readouts of urinary DNA barcodes are used to classify lung cancer groups from healthy cohorts. **(B)** For *in vivo* validation of inhalational delivery of the ABNs, a nose-only exposure inhalation tower was assembled with a vibrating-mesh nebulizer at the bottom and the mice situated in restrainers on the side. The aerosolized ABNs flow into the inhalation tower with medical airs. The mice were only exposed to the ABNs-laden aerosols only via nasal openings. The median volumetric diameter of generated aerosols was in the range of 2-4 µm.

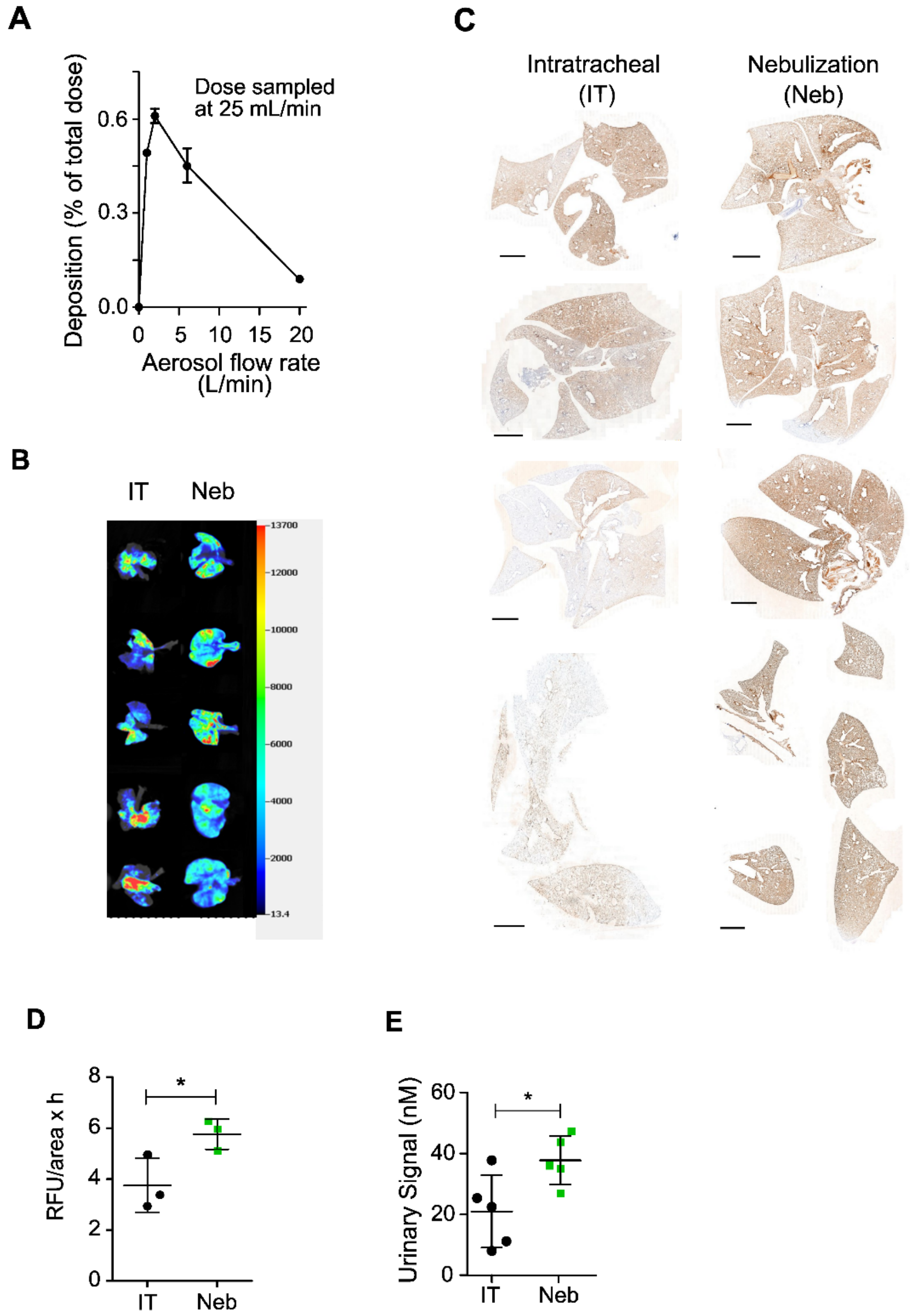


**Figure S4. Local biodistribution of inhaled and intratracheally injected ABNs in lungs. (A)** Compressed air flow rate was optimized to maximize ABN aerosols available for animals to inhale. Nebulization outperforming intratracheal injection enables broad and even distribution of ABN in lungs as revealed by **(B)** fluorescence tracers and **(C)** immunohistochemistry staining. Biotin-labeled ABN were delivered to wild-type C57BL/6 mice and the lungs excised 20 minutes after administration from the intratracheally injected (IT) and nebulization (Neb) cohorts (n=4) were fixed overnight by 4% paraformaldehyde. The fixed lungs were then stained with streptavidin-HRP. The scale bar is 2mm. **(D)** plasma concentrations and **(E)** urinary readouts of cleaved reporters in intratracheal and nebulization cohorts. Nebulization groups showed higher area under curve (AUC) of pharmacokinetics and urinary reporter concentrations. Abbreviation: RFU = relative fluorescence units.

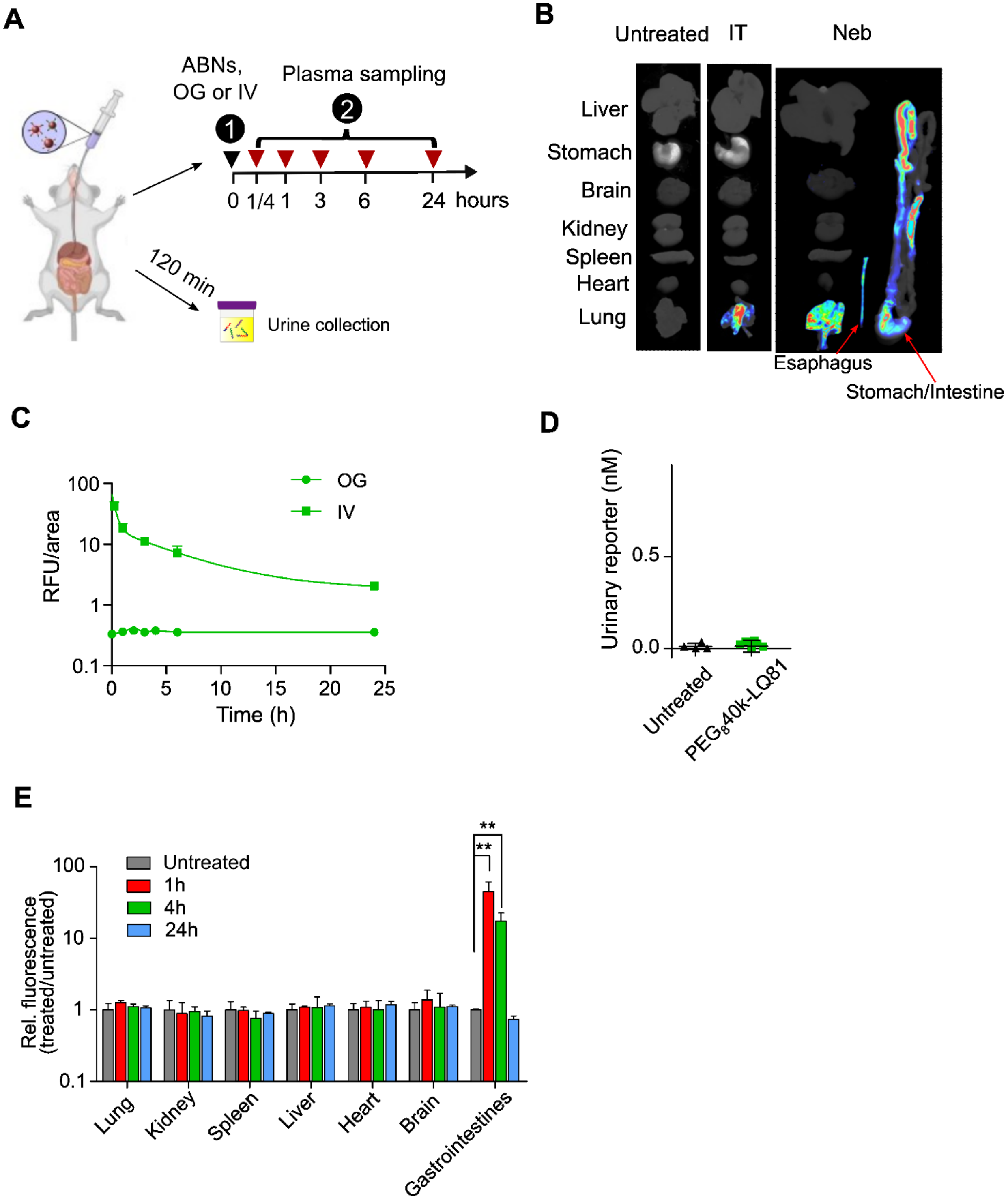


**Figure S5. Orally administered ABNs pose no interference with urinary readouts. (A)** Schematic illustration of the delivery of Cy7-tagged ABNs with Glu-fib reporters to animals via oral gavage (OG). The plasmas were sampled at various time points to monitor the absorption of ABN into systemic circulation. The urines were collected at 120 min post oral gavage and the fluorescence of Cy7 in urine samples was measured. The biodistribution was examined2 min, 0.25h, 1 h, 4h, and 24 h post oral administration. **(B)** Representative 1 h biodistribution of intratracheally-instilled (IT) and nebulized (Neb) ABNs that were tagged with Cy7. The ABNs via IT were solely distributed in lungs with heterogeneous patterns, whereas nebulized ABNs were inhaled into not only the lungs, but digestive system, including esophagus. **(C)** Plasma concentration of Cy7 that summed up ABNs and Cy7-tagged reporters liberated from ABNs. Fluorescent signals were not detected in the plasma of the mice treated with orally delivered ABNs, indicating no ABNs or cleaved reporters can re-enter the systemic circulation that ultimately led to urinary concentration. Intravenously injected ABNs (IV) was used as a positive control. **(D)** Urinary readouts of Cy7 fluorescence in the mouse group of oral gavage shows no significant difference in comparison with untreated counterparts, corroborating that orally administered ABNs fail to release DNA reporters to urines. **(E)** Systemic biodistribution of ABNs at 1, 4 and 24 hours post oral administration. No obvious observation of ABNs in tissues outside the digestive system suggests oral ABNs were excreted without no entry into systemic circulation. Cy7 fluorescence was not detected in the kidneys of mice dosed orally with ABNs, further corroborating undetectable urinary readouts in (D).

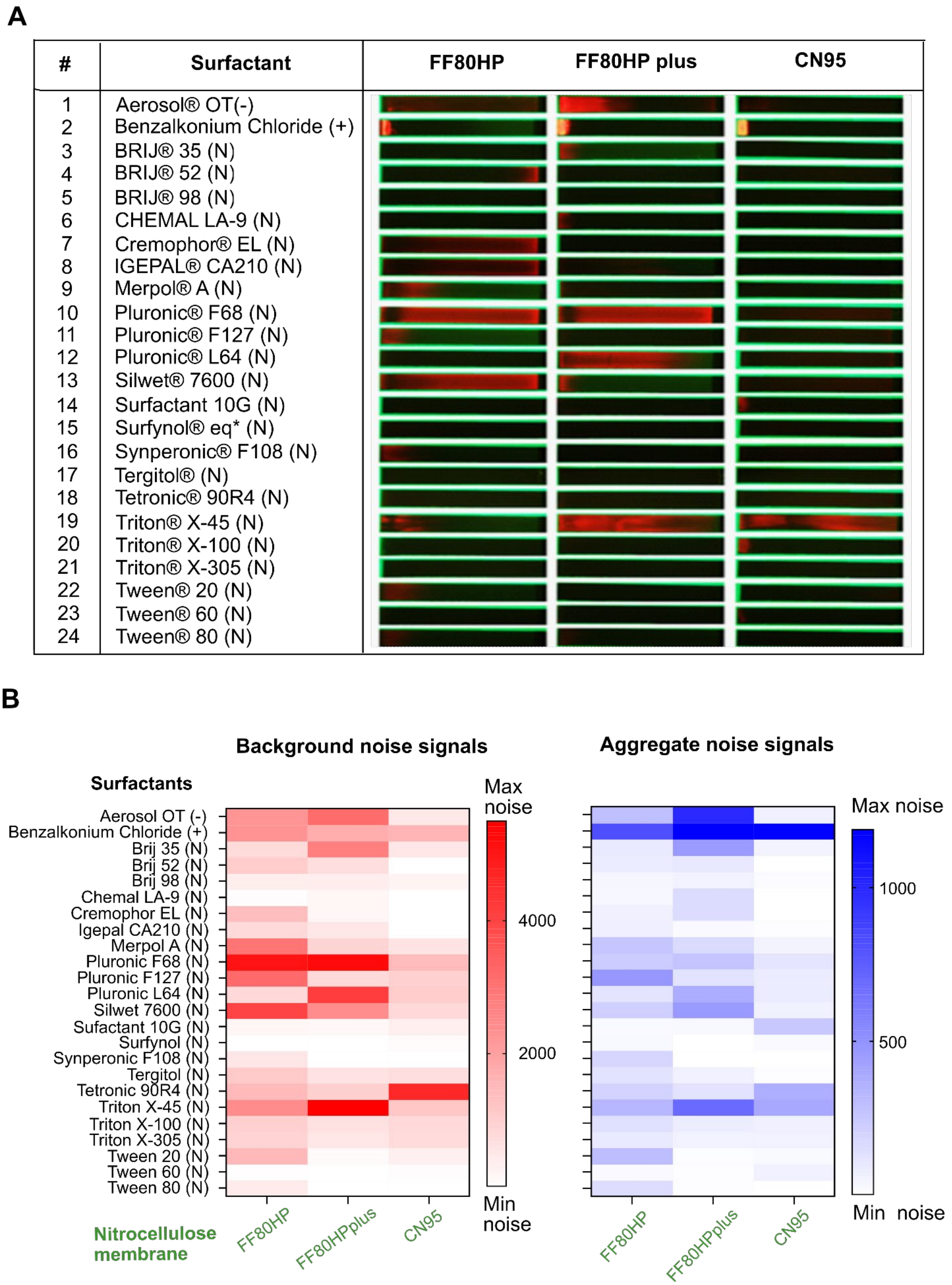

A
#
Surfactant
FF80HP
FF80HP plus
CN95
1 Aerosol® OT(-)
2 Benzalkonium Chloride (+)
3 BRIJ® 35 (N)
4 BRIJ® 52 (N)
5 BRIJ® 98 (N)
6 CHEMAL LA-9 (N)
7 Cremophor® EL (N)
8 IGEPAL® CA210 (N)
9 Merpol® A (N)
10 Pluronic® F68 (N)
11 Pluronic® F127 (N)
12 Pluronic® L64 (N)
13 Silwet® 7600 (N)
14 Surfactant 10G (N)
15 Surfynol® eq* (N)
16 Synperonic® F108 (N)
17 Tergitol® (N)
18 Tetronic® 90R4 (N)
19 Triton® X-45 (N)
20 Triton® X-100 (N)
21 Triton® X-305 (N)
22 Tween® 20 (N)
23 Tween® 60 (N)
24 Tween® 80 (N)
B
Background noise signals
Aggregate noise signals
Surfactants
Aerosol OT (-)
Benzalkonium Chloride (+)
Brij 35 (N)
Brij 52 (N)
Brij 98 (N)
Chemal LA-9 (N)
Cremophor EL (N)
Igepal CA210 (N)
Merpol A (N)
Pluronic F68 (N)
Pluronic F127 (N)
Pluronic L64 (N)
Silwet 7600 (N)
Sufactant 10G (N)
Surfynol (N)
Synperonic F108 (N)
Tergitol (N)
Tetronic 90R4 (N)
Triton X-45 (N)
Triton X-100 (N)
Triton X-305 (N)
Tween 20 (N)
Tween 60 (N)
Tween 80 (N)
Max noise
4000
2000
Min noise
Max noise
1000
500
Min noise
Nitrocellulose membrane
FF80HP
FF80HPplus
CN95
FF80HP
FF80HPplus
CN95

**Figure S6.Screening of surfactants in running buffer against commercially available membrane. (A)** Surf's Up® Surfactant Kit containing up to 24 different surfactants were used for lateral flow assay development. The kit mainly covered non-ionic surfactants by also contained 1 cationic and 1 anionic surfactant. Running buffer prepared using these surfactants were tested against 3 commercially available membranes, namely FF80HP, FF80HP Plus and CN95. Membranes were dipped into the running buffer solution (1% surfactants and 0.5% Eu-conjugates) and the fluorescent signal was recorded. Candidates that 1) did not result in conjugate aggregation at the immersed section of the strip and 2) did not cause high background fluorescence were considered for further development. In other words, strips with minimum fluorescence across the membrane were nominated. * denotes 2,4,7,9-tetramethyl-5-decyne-4,7-diol ethoxylate, which is used as a Surfynol® equivalent. (+) denotes cationic properties, (-) denotes anionic properties, and (N) denotes non-ionic properties. **(B)** Tabulated heat maps indicate the quantification of background noise signals (e.g. surfactant 10 Pluronic F68 in **(A)**), implying non-specific binding, and conjugate aggregation signals (e.g. surfactant 2 Benzalkonium chloride in **(A)**) at the immersed section of the strip, respectively. To generate the heatmap of background noise signals, Eu fluorescence intensity (illustrated in **Fig.5C**) profile was first plotted with respect to the distance from the left on the strips using ImageJ. Area under the curve (AUC) of the profile was obtained with GraphPad Prism 5.0. The AUC values were evaluated as background noises (due to non-specific binding). High AUC values indicate high non-specific binding on the strips. Regarding the heatmap of conjugate aggregation, we delineated the first 2.1 mm from the left on the strip as the immersion onset and measured the AUC of Eu intensity profiles in this part. We can observe Tween 60 has one of lowest AUC in both heatmaps (especially), while other surfactants are high in at least one heatmap.

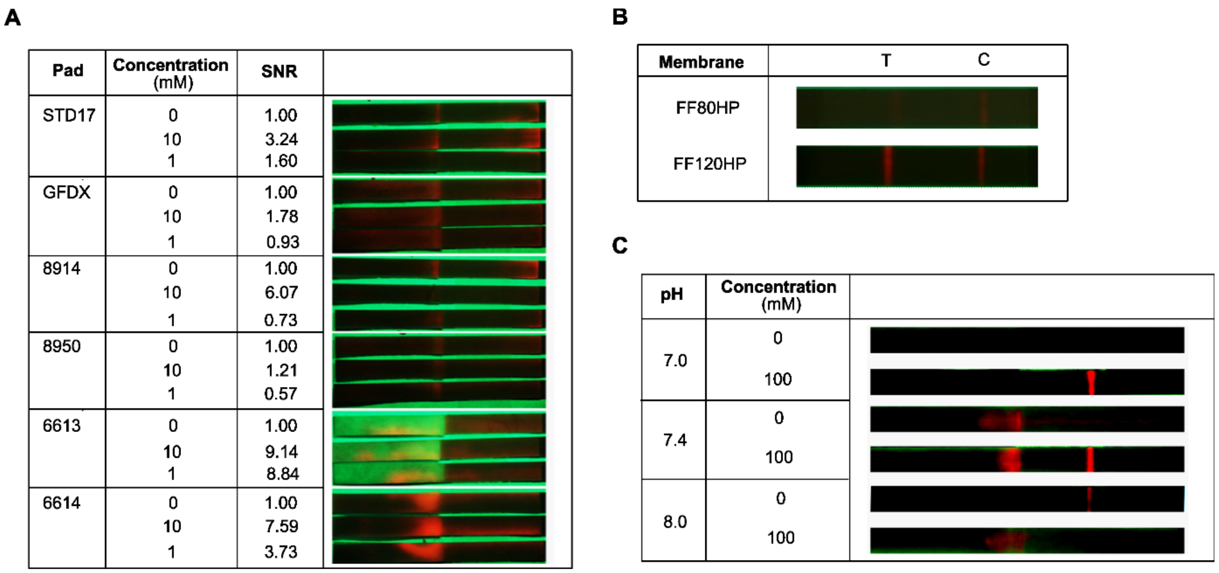


**Figure S7. Optimization of lateral flow assay (A)** Screening of conjugate pads for the optimal release of Eu-conjugates. Commercially available conjugate pads containing Eu conjugates were laminated onto FF80HP Plus immobilized with capture probe for barcode 1. Urine containing 0, 1 and 10 nM of the barcode were spotted onto the conjugate pads. Strips were immersed into running buffer of pH 8.6 containing 100 mM Borate, 150 mM NaCl and 10% Tween 60. Criteria to determine optimal performance of the pad included the efficient release of the conjugates (minimal aggregation at the junction between the pads), low background signal and high signal to noise ratio of the barcode. Although the fluorescent signal was not clear visually on the test line, it can be quantified using reader systems. Based on the results, conjugate pad STD 17, 8914 and 8950 were selected for further development. **(B)** Further development of the performance of nitrocellulose membrane. The use of FF120HP Plus with Pluronic F68 showed a more sensitive detection of barcode 2 in comparison to FF80HP previously optimized. Abbreviation: C=control line, T=testing line. **(C)** Optimization of running buffer pH with casein pre-treated Eu-Neutravidin. Casein treatment further resulted in the reduction of background signal. pH of the running buffer played a role in the performance of the barcode quantification.

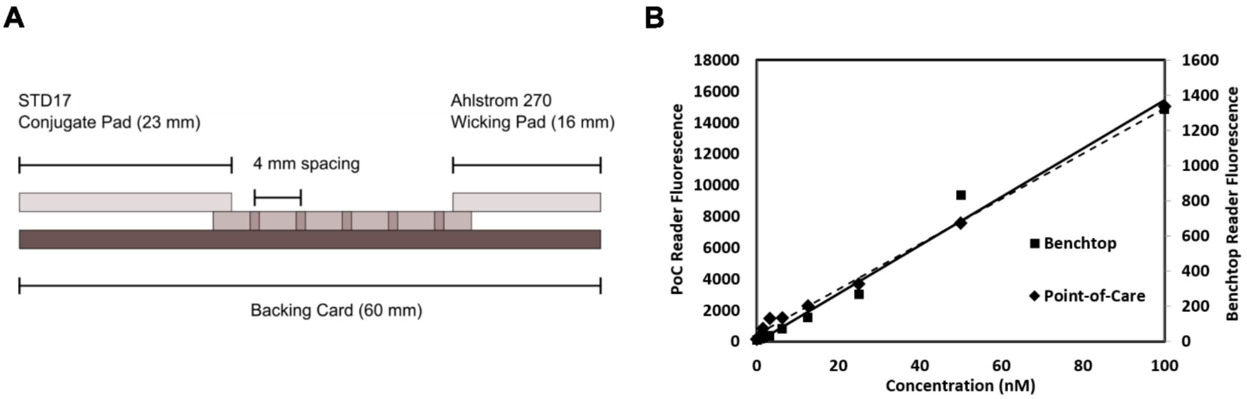


**Figure S8. (A)** Dimensions of the multiplexed lateral flow assay developed in this study. **(B)** Performance of point-of-care reader vs benchtop reader. Both readers show similar performance in the range of interests.

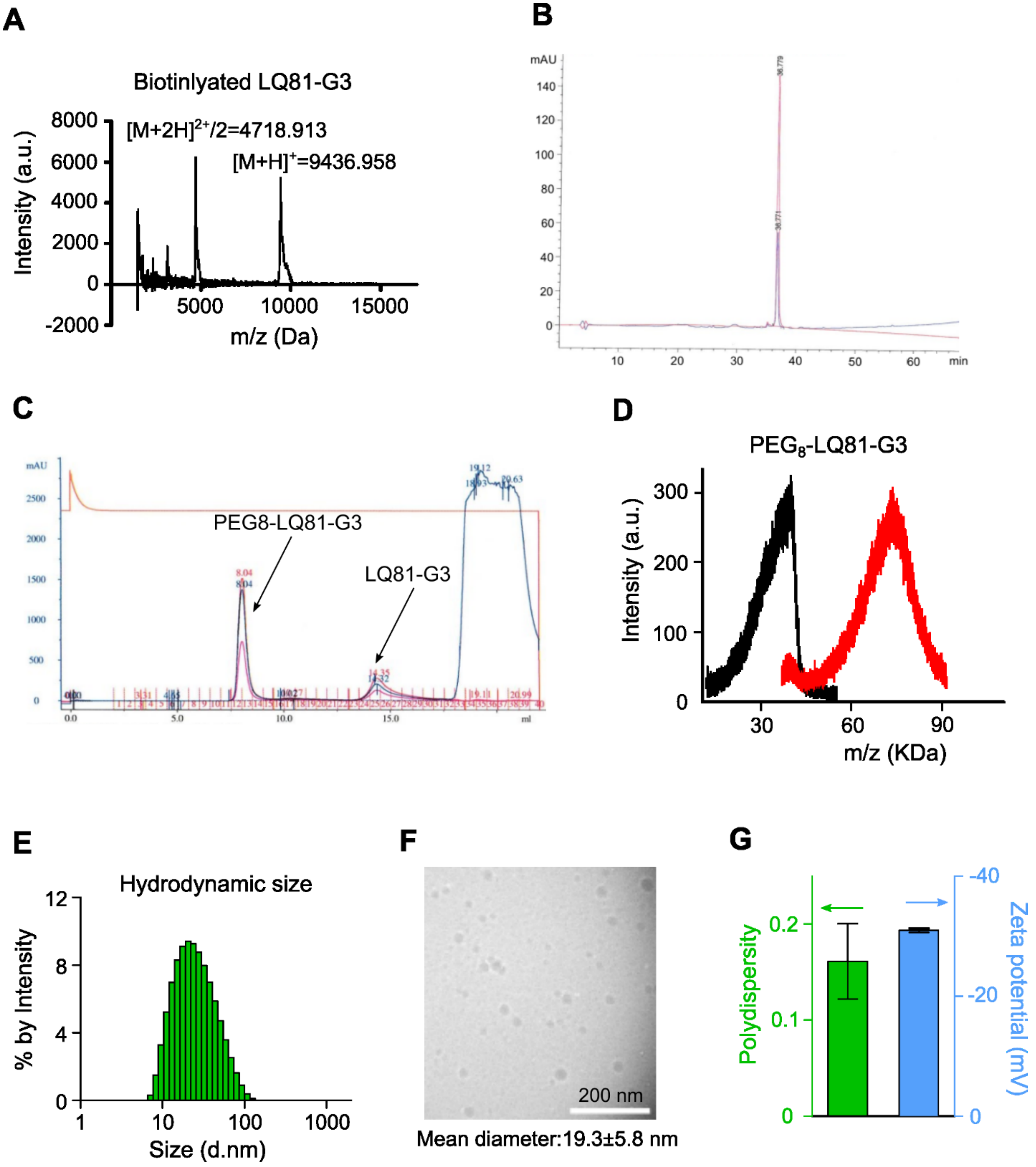


**Figure S9. Synthesis and characterization of activity-based nanosensors with synthetic DNA reporters. (A)** Mass spectrum of the LQ81-DNA3 conjugate. **(B)** Biotinylated LQ81-DNA3 conjugate was purified with a HPLC equipped with a reverse-phase C18 column to remove unreacted DNA barcodes and peptide LQ81. The eluent composed of acetonitrile and water gradually ramped up the ratio from 10/90 to 90/10 in a period of 30 min. **(C)** $PEG_8$40k-LQ81-DNA3 conjugate was purified with fast protein liquid chromatography (FPLC) with a Superdex 75 10/300 GL column. The conjugate was eluted out at 8.04 min, whereas the unreacted LQ81-DNA3 conjugate was eluted from the column at approximately 14.3 minutes. **(D)** Mass spectrum of the $PEG_8$40k-LQ81-DNA3 conjugate, indicating approximately 4 DNA barcodes were conjugated to each PEG scaffold. Black: 8-arm PEG-maleimide, red:

$PEG_8$40k-LQ81-DNA3 conjugate. **(E)** Dynamic light scattering (DLS) showed hydrodynamic size of mass-coded, 8-arm PEG ABNs is approximately 18 nm. The measurement was performed in 1X PBS. **(F)** Cryo-transmission electron micrograph (Cyro-TEM) of the DNA-coded nanosensors. Geometric size calculated from the Cryo-TEM was 20 nm on average (calculated based on n=50 particles). **(G)** Polydispersity index of 8-arm PEG ABNs measured by DLS was 0.16 and DNA barcodes confer the PEG nanoscaffold negative charges upon conjugation.

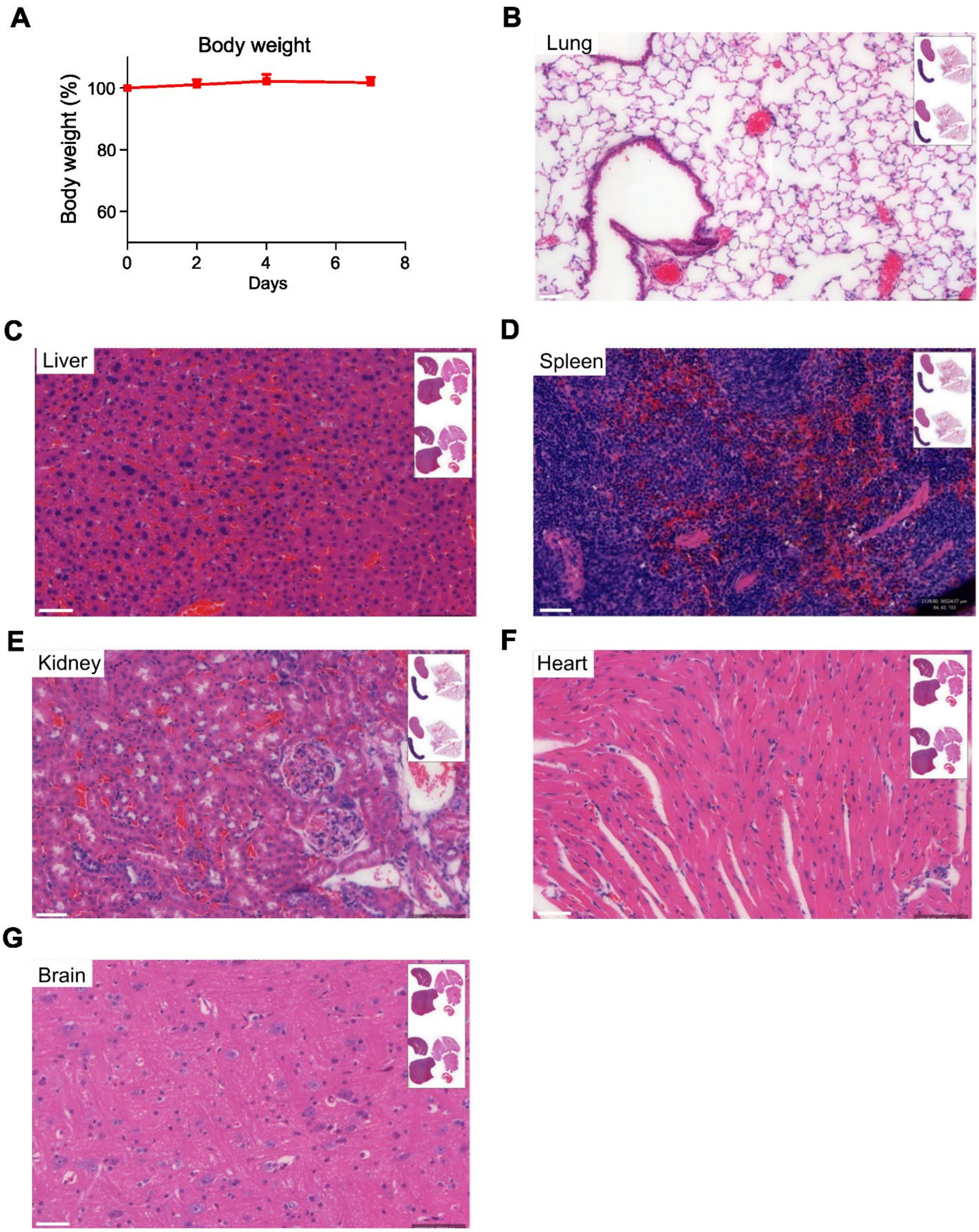


**Figure S10. No obvious toxicity was observed in mice treated with nebulized nano-in-micro sensors 7 days post administration**. **(A)** Body weights of 5 mice remained largely unchanged over 7 days post nebulization of a single dose of 4-plex ABNs that is 3 folds of the dose used for detection. Representative H&E histological images of excised

**(B)** lung, **(C)** liver, **(D)** spleen, **(E)** kidney, **(F)** heart, and **(G)** brain. Scale bar is 50 μm long. All these results demonstrated no detectable acute general toxicity was observed associated with inhaled DNA-coded ABNs.

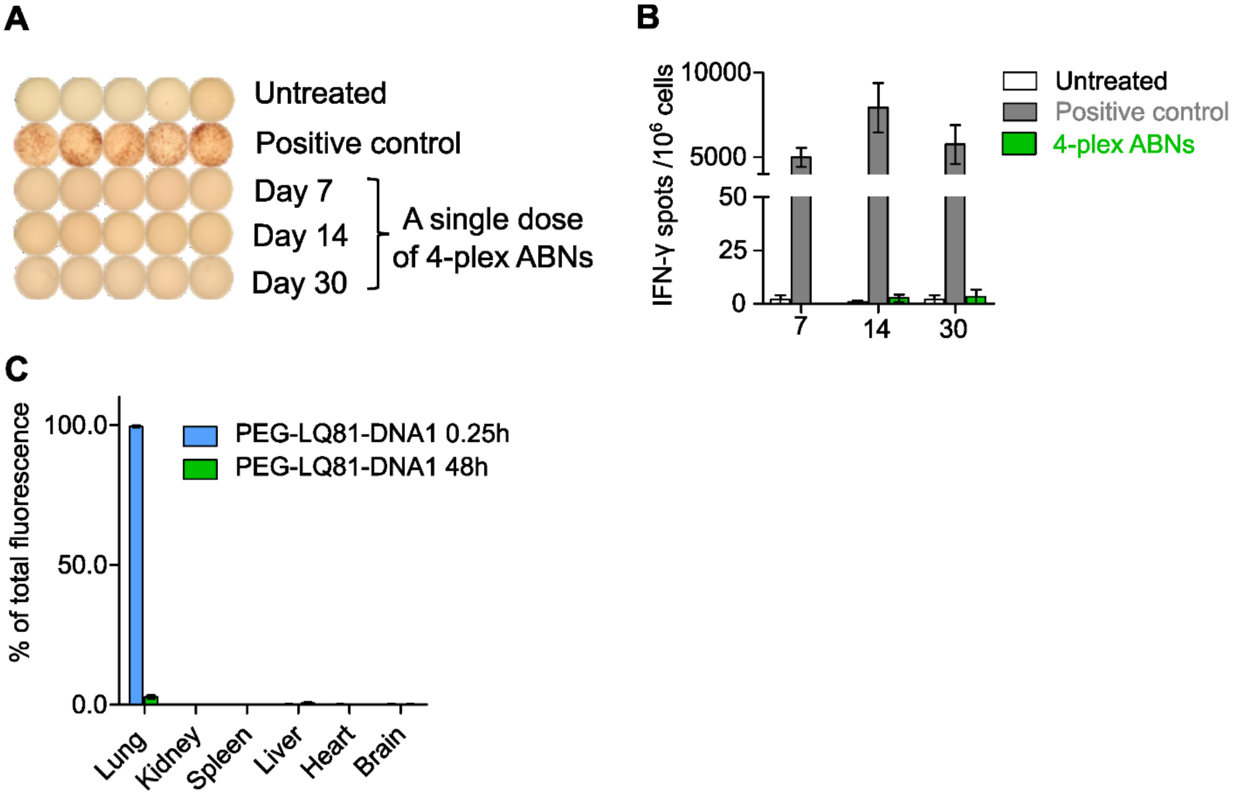


**Figure S11. Four-plex ABNs with DNA barcodes were non-immunogenic at the delivered dose. (A)** Representative images of IFN-γ ELISPOT assay of untreated, stimulated (positive control) and ABN-treated peripheral white blood cells. In the positive control, the white blood cells were stimulated with… **(B)** Quantification of IFN-γ spots of untreated and treated samples revealed non-immunogenicity of a single dose of 4-plex ABNs that is 3 folds of the dose used for detection. **(C)** Biodistribution (48h) of DNA-barcoded ABNs delivered via nebulization. In control group (0.25h), the mice were euthanized 0.25 h post ABN nebulization and organs were then harvested to quantify fluorescence from the ABN.

**Table S1. Panel of prominent peptide sequences responsive to a library of 21 proteases upregulated in human Stage I LUAD.** Uppercase letters denote L-isomers of amino acids, lowercase letters denote D-isomers of amino acids. Abbreviation: GzmB, NE and CatK denote a peptide sequence is cleaved by human granzyme B, neutrophil elastase and cathepsin K, respectively.

| **Peptide code** | **Sequences (C terminus= $CONH_2$)** |
|---|---|
| **LQ2** | GGRQARAVGGC |
| **LQ3** | GGRRARVVGGC |
| **LQ28** | GGLAQAFRSGC |
| **LQ31** | GGRQRRALEKGC |
| **LQ35** | GGPRPFNYLGC |
| **LQ38** | GGAKIRGQAKGC |
| **LQ46** | GGSGDRMWggGC |
| **LQ49** | GGKLRVVGGHPGC |
| **LQ54** | GGNIPMGLLYNKGC |
| **LQ70** | GGLGALLRVKRLEC |
| **LQ79** | GGPQGIWGQGC |
| **LQ81** | GGPVPLSLVMC |
| **LQ83** | GGfPRSGGGC |
| **LQ86** | GGPLGMRGGC |
| **LQ87** | GGAPFEMSAGC |
| **P2** | GGPVGLIGC |
| **P13** | GGP-(Cha)-G-Cys(Me)-HAGC |
| **GzmB** | GGAIEFDSGC |
| **NE** | GGAAFAGC |
| **CatK** | GGHPGGPQC |

**Table S2. Panel of 20-plexed ABNs.** The unique mass-coded glutamic acid-labeled fibroprotein was chemical bonded to each peptide sequence in Table 1 via a photocleavable linker. The entire peptides were then conjugated to 8-arm PEG nanoscaffolds via maleimide-thiol click chemistry. The ABNs were named by adding a prefix ‘P’ to peptide codes, indicating the PEG nanocore. . Uppercase letters denote L-isomers of amino acids, lowercase letters denote D-isomers of amino acids. The ABNs were purified with HPLC and molecular weights were determined with MALDI-TOF.

| ABN code | Sequence | Molecular weight (Da) |
|---|---|---|
| **PLQ2** | e(+3G)(+1V)ndneeGFFs(+4A)r-ANP-GGRQARAVGGC-(PEG8-40kDa) | 65224.0 |
| **PLQ3** | e(+2G)Vndnee(+2G)FFs(+4A)r-(ANP)-GGRRARVVGGC-(PEG8-40kDa) | 64035.2 |
| **PLQ28** | eGVndnee(+3G)(+1F)Fs(+4A)r-(ANP)-GGLAQAFRSGC-(PEG8-40kDa) | 64640.8 |
| **PLQ31** | e(+2G)(+6V)ndnee(+3G)(+1F)(+1F)s(+1A)r-(ANP)-GGRQRRALEKGC-(PEG8-40kDa) | 64769.9 |
| **PLQ35** | e(+3G)(+1V)ndneeG(+10F)FsAr-(ANP)-GGPRPFNYLGC-(PEG8-40kDa) | 66570.5 |
| **PLQ38** | e(+2G)Vndnee(+2G)F(+10F)sAr-(ANP)-GGAKIRGQAKGC-(PEG8-40kDa) | 63514.2 |
| **PLQ46** | eGVndneeGF(+10F)s(+4A)r-(ANP)-GGSGDRMWggGC-(PEG8-40kDa) | 66000.5 |
| **PLQ49** | e(+2G)(+6V)ndneeG(+10F)(+1F)s(+1A)r-(ANP)-GGKLRVVGGHPGC-(PEG8-40kDa) | 65018.9 |
| **PLQ54** | eG(+6V)ndneeG(+10F)Fs(+4A)r-(ANP)-GGNIPMGLLYNKGC-(PEG8-40kDa) | 67839.9 |
| **PLQ70** | e(+3G)(+1V)ndnee(+2G)(+10F)Fs(+4A)r-(ANP)-GGLGALLRVKRLEC-(PEG8-40kDa) | 66952.9 |
| **PLQ79** | e(+2G)Vndnee(+3G)(+10F)(+1F)s(+4A)r-(ANP)-GGPQGIWGQGC-(PEG8-40kDa) | 66158.7 |
| **PLQ81** | eGVndneeG(+10F)(+10F)sAr-(ANP)-GGPVPLSLVMC-(PEG8-40kDa) | 66457.6 |
| **PLQ83** | eG(+6V)ndnee(+3G)(+1F)Fs(+4A)r-(ANP)-GGfPRSGGGC-(PEG8-40kDa) | 60210.3 |
| **PLQ86** | e(+2G)VndneeG(+10F)(+10F)s(+4A)r-(ANP)-GGPLGMRGGC-(PEG8-40kDa) | 63497.7 |
| **PLQ87** | e(+2G)(+6V)ndnee(+3G)(+10F)(+1F)s(+4A)r-(ANP)-GGAPFEMSAGC-(PEG8-40kDa) | 64116.1 |
| **PP2** | e(+3G)(+1V)ndnee(+2G)(+10F)(+10F)sAr-(ANP)-GGPVGLIGC-(PEG8-40kDa) | 60463.1 |
| **PP13** | eGVndnee(+2G)(+10F)(+10F)s(+4A)r-ANP-GGP-(Cha)-G-Cys(Me)-HAGC-(PEG8-40kDa) | 60949.4 |
| **PGzmB** | e(+2G)(+6V)ndneeGFFsAr-(ANP)-GGAIEFDSGC-(PEG8-40kDa) | 59710.9 |
| **PNE** | eG(+6V)ndneeG(+10F)(+10F)sAr-(ANP)-GGAAFAGC-(PEG8-40kDa) | 59565.2 |
| **PCatK** | eG(+6V)ndneeGF(+1F)s(+1A)r-(ANP)-GGHPGGPQC-(PEG8-40kDa) | 61364.2 |

**Table S3. DNA sequences of barcodes and capture probes in the study**

| Barcode | Sequence |
|---|---|
| **DNA1** | /5BiosG/G*T*T*A*G*T*A*A*G*A*A*C*T*A*T*T*T*G*A*A/3DBCON/ |
| **DNA2** | /5BiosG/T*G*T*C*T*A*T*A*A*A*A*C*A*T*T*A*A*G*A*T/3DBCON/ |
| **DNA3** | /5BiosG/T*C*A*T*A*G*T*T*A*T*C*T*T*A*A*C*A*A*T*C/3DBCON/ |
| **DNA4** | /5BiosG/T*C*A*T*A*G*T*T*A*G*C*G*T*A*A*C*G*A*T*C/3DBCON/ |
| **C1** | /5AmMC6/TTCAAATAGTTCTTACTAAC |
| **C2** | /5AmMC6/ATCTTAATGTTTTATAGACA |
| **C3** | /5AmMC6/GATTGTTAAGATAACTATGA |
| **C4** | /5AmMC6/GATCGTTACGCTAACTATGA |

*denotes phosphorothioate bond

/5BiosG/, /3DBCON/ and /5AmMC6/ are the modification code using IDT synthesis service

**Acknowledgments:** We thank A. Mancino from Syneos Health for performing mass spectrometry and DCN Technologies for optimizing and manufacturing paper strips for lateral flow assay. We are also grateful to the Koch Institute Swanson Biotechnology Center, specifically the Histology core, the Biopolymer and Proteomics core, and Nanotechnology Materials core. **Funding**: This work was supported by Johnson & Johnson Lung Cancer Initiative and Howard Hughes Medical Institute (to S.N.B.)., and partly by a Koch Institute Support (core) Grant P30-CA14051 from the National Cancer Institute. S.N.B. is an HHMI investigator. **Author contributions**: Q.Z., E.K.W.T., M.C.M and S.N.B. conceived and designed the study. Q.Z., E.K.W.T. and M.C.M. performed experiments. Q.Z. and E.K.W.T performed all statistical analysis. Q.Z. and T.P. generated animal models. Q.Z., E.K.W.T., M.C.M, L.H., T.F., H.E.F. and S.N.B. supervised the research. Q.Z., E.K.W.T., and S.N.B. wrote the first draft of the manuscript. H.E.F. assisted in the preparation of the manuscript. All authors contributed to writing and editing subsequent drafts of the manuscript and approved the final manuscript. **Competing interests**: T.J. is a member of the Board of Directors of Amgen and Thermo Fisher Scientific and a co-founder of Dragonfly Therapeutics and T2 Biosystems; serves on the Scientific Advisory Board of Dragonfly Therapeutics, SQZ Biotech, and Skyhawk Therapeutics; and is President of Break Through Cancer. T.J.'s laboratory currently receives funding from Johnson & Johnson and The Lustgarten Foundation, but these funds did not support the research described in this manuscript. S.N.B. reports compensation for cofounding, consulting for, and/or board membership in Glympse Bio, Satellite Bio, CEND Therapeutics, Catalio Capital, Intergalactic Therapeutics, Port Therapeutics, Vertex Pharmaceuticals, and Moderna, and receives sponsored research funding from Johnson & Johnson, Revitope, Owlstone and Howard Hughes Medical Institute. The remaining authors declare no competing interests. **Data and materials availability**: All data associated with this study are present in the paper and Supplementary Materials.
